# Convergent and divergent spatial topographies of individualized brain functional networks and their developmental origins

**DOI:** 10.1101/2025.11.18.689167

**Authors:** Jianlong Zhao, Yu Zhai, Yuehua Xu, Lianglong Sun, Tengda Zhao

## Abstract

The human brain is intrinsically organized into canonical functional networks with distinct spatial topographies. While precision functional mapping delineates individualized topographies of single networks, the spatial coordination among networks and its developmental origin remains largely unknown. Here, we propose functional topography covariance analysis (FOCA), a novel framework that quantifies convergent and divergent spatial alignments across individualized functional networks and further delineates their internetwork relationships, neurobiological basis, ontogenetic layouts, and cognitive outcomes. Across two well-established task-free functional MRI (fMRI) datasets encompassing both conventional and densely sampled scans, FOCA consistently revealed self-clustered hierarchies in the coordination of functional topographies, closely aligned with existing functional gradients and characterized by convergent couplings within primary systems and divergent couplings in higher-order systems. Such pattern was well predicted by fundamental neurobiological attributes, especially aerobic glycolysis. In a large public neonatal cohort, FOCA matrix exhibited adult-inverted hierarchical couplings and marked changes in auditory and action-mode networks, primarily driven by prominent redistributions of negative couplings. Moreover, neonatal FOCA profiles in the primary visual system significantly predicted neurodevelopmental outcomes at 18 months. Finally, compared with conventional functional connectivity (FC), FOCA proved more robust to global signals and more sensitive to the maturation of negative couplings. These findings highlight the critical role of negative connectivity in brain functional reorganization and advance our understanding of the cooperative and competitive relationships among functional systems and their developmental origins.

## Introduction

The human brain achieves complex functional coordination through continuously cooperative and competitive neuronal activation ^[1–3]^. This process manifests as the emergence of canonical functional networks ^[4, 5]^ at the macroscale with unique brain locations, highly individual-specific boundaries and diverse connectivity profiles ^[6–8]^. These spatial architectures of functional networks, conceptualized as functional topography, collectively reflects how neuronal circuits are spatially activated and interact across the cortex, providing a comprehensive characterization of the inherent brain functional organization ^[7, 9]^. The refinements of functional topographies during development also closely encode the maturation of the brain and behaviour ^[10, 11]^. However, existing studies on brain functional topography mainly focused on the spatial configuration of single networks, and the spatial coordination between networks and their developmental origin remain largely unknown.

Functional connectome modelling enables the systematic estimation of functional interactions between brain areas ^[12, 13]^. Typically, network nodes are defined by cortical parcellations derived from atlas-based priors, and edges are characterized by temporally correlated BOLD fluctuations ^[4, 5]^. However, these approaches involve individual functional parcellations through structural registrations to a standard atlas, which are unable to capture individualized neural activity patterns that deviate from those of common structural scaffolds ^[7, 8]^. Moreover, edges with negative correlations are generally considered unreliable and excluded from the network ^[14, 15]^, which implies that valuable anticorrelated relationships between function parcels are largely ignored. Neurophysiological evidence has shown that inhibitory neuronal activity is abundant in cortical microcircuits and plays crucial roles in generating functionally specialized neural oscillations, suppressing noise interference, and supporting synaptic plasticity ^[16, 17]^. A reliable estimation of antagonistic interactions between large-scale functional networks is critical for understanding the functional segregation and functional flexibility of the brain ^[1, 7]^. Recent advances in individualized functional network mapping using highly sampled resting-state fMRI scans have enabled the identification of personal trait-like functional networks ^[18, 19]^. The spatial topographies of these intrinsic function parcels exhibit substantial variability in both location and shape across individuals but are highly consistent within individuals ^[18, 20]^. Guided by prior cortical seeds, task-like functional activation patterns could also be derived using resting-state scans ^[21–23]^ and show high behaviour and neurobiological relevance ^[6, 10]^. Emerging studies emphasize that the spatial topography of personalized functional networks also affects their internetwork relationships ^[1]^. For instance, the action-mode network (AMN) and the default mode network (DMN) exhibit quite complementary spatial topographies with strongly anticorrelated fluctuation patterns, reflecting the balance between the default-mode state and goal-directed behavioural processes. However, the systematic characterization of internetwork interactions between individualized functional networks and their neurobiological foundations remains largely unaddressed.

A fundamental goal of modern neuroscience is to elucidate how brain functional architecture originates and develops ^[24–26]^. Recent fMRI studies on brain development have predominantly used personalized network modelling approaches rather than group-level frameworks, to reduce the methodological biases when neglecting the variation in the functional boundaries among individuals ^[10, 26–28]^. Such studies have revealed that the individualized functional topography is refined dramatically with brain maturation at pivotal developmental stages. In neonates, major functional networks have already become identifiable, and the adjustments are concentrated in visual and motor networks ^[29]^. In adolescents, boundaries of the frontoparietal network are refined prominently during development and significantly predict individual executive function ^[10, 11]^. However, previous studies have focused mainly on age-related changes in the functional topography of single networks. Characterizing how functional networks spatially coordinate with each other across development is crucial for understanding the integration of and segregation between functional systems during brain maturation.

To fill these gaps, we proposed functional topography covariance (FOCA) analysis, a novel framework that quantifies the internetwork coupling of individualized functional networks by estimating the covariance among their spatial topography maps. Leveraging three well-established task-free fMRI datasets encompassing conventional and densely sampled scans, across neonatal and adult cohorts ^[7, 30, 31]^, we systematically investigated the interplay between individualized functional networks and their developmental origins. First, we generated a canonical adult representation of FOCA and describe its individual variability pattern. Population consistency and scan duration stability were also evaluated. Then, we resolved the hierarchical organization of the FOCA matrix and delineated its neurobiological underpinnings using machine learning models on sets of brain annotation atlases. Next, leveraging a large sample of neonatal scans, we quantified how neonatal FOCA maps differ from those of adults and whether they can predict brain maturation at early postnatal period and neurodevelopmental outcomes at 18 months. Finally, we compared FOCA with conventional functional connectivity (FC) with respect to the methodological stability and sensitivity in capturing negative connections.

## Results

### Data samples

The Midnight Scan Club (MSC) dataset ^[7]^ (N = 9, 300 minutes scans) provided dense sampled adult brain scans, enabling the initial mapping of the typical FOCA matrix. The Human Connectome Project (HCP) dataset ^[31]^ (N = 997, 58 minutes scans) was used to validate the FOCA matrix in a large adult cohort with relatively short scans and to measure the individual variability of FOCA. Finally, the Developing Human Connectome Project (dHCP) dataset ^[30]^ (N = 422, 15 minutes scans; Fig. S1), a large public neonatal fMRI cohort, was employed to delineate the typical neonatal FOCA map, assess differences from the map of adults, chart early postnatal trajectories, and predict neurodevelopmental outcomes at 18 months.

### Individualized spatial topography of positive and negative connections in adult functional networks

We leveraged a widely used template-matching (TM) approach ^[27, 32, 33]^ to generate individualized functional networks. A recently proposed fine-grained atlas containing 11 major functional networks with 20 subnetworks (for details on each network, see Fig. S2) was adopted to obtain the initial group-level constraint ^[7, 34–37]^. Individualized networks were generated by constructing customized network templates (Fig. S3) and matching the individual connectivity profile of each vertex to the template (Fig. S4). This procedure proved highly effective in the MSC dataset, as the individualized networks exhibited significantly higher within-network homogeneity than the group-level parcellations (t = 6.23, *P* = 1.20 × 10⁻^5^, Cohen’s d = 5.69; Fig. S5). Then, we derived individualized topographic maps by correlating the mean time series of each network with all cortical vertices (Fig. S4). Topographic maps for all eleven functional networks are shown in Figure 1. Visual inspection revealed that the coarse spatial topography of both positive and negative functional connectivity was largely consistent across individuals (Fig. 1). At a fine spatial scale, substantial inter-individual differences emerged. As an illustration, the DMN consistently showed positive connectivity in the medial prefrontal and precuneus cortices and negative connectivity in the dorsal anterior cingulate (dACC) and anterior insula cortices. Yet, the precise location and spatial extent of these connections varied substantially across individuals. Comparable patterns were also observed across the remaining functional networks.

**Fig. 1.**
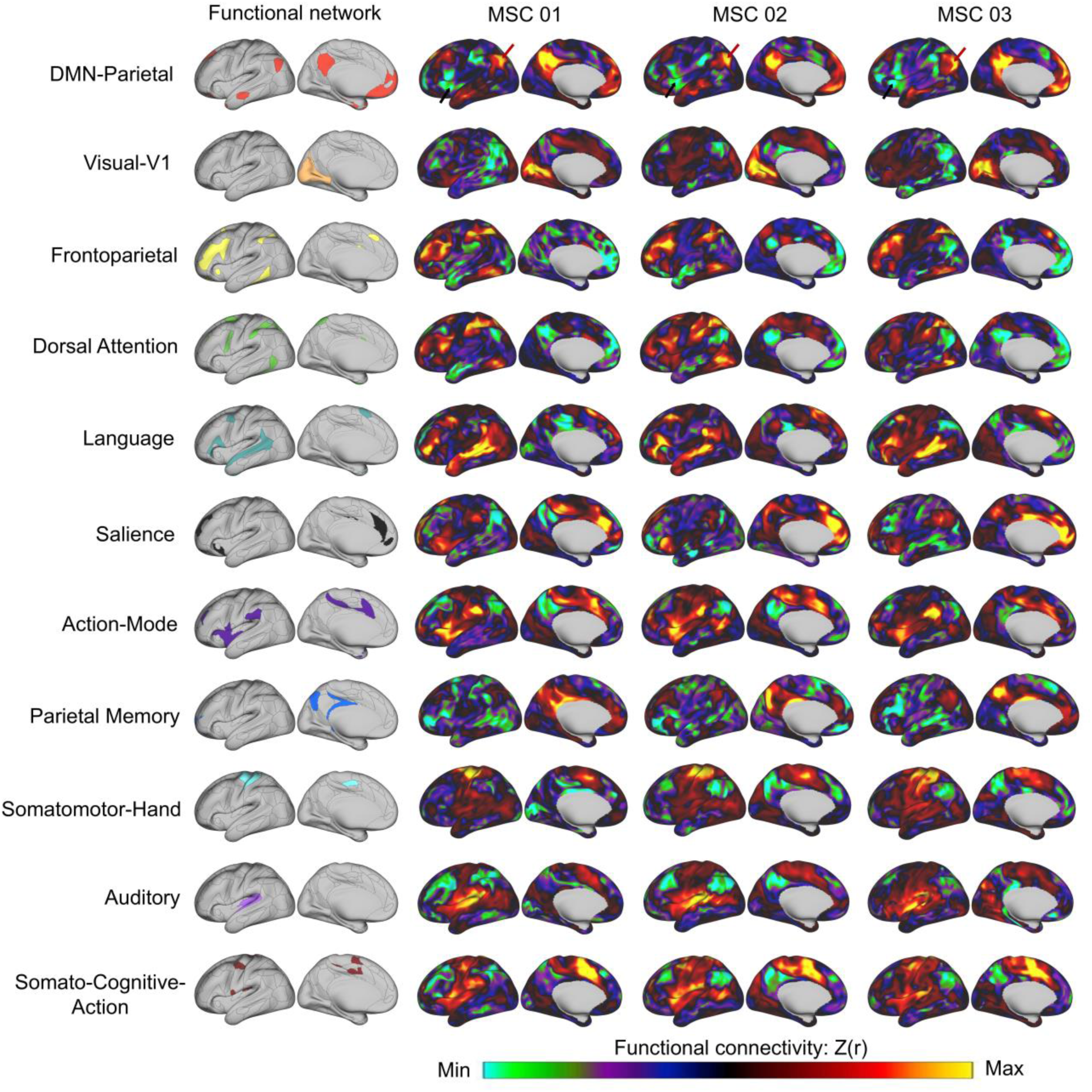
Participant-specific functional topography maps derived from individualized functional networks. The left column shows representative seed networks, and the right columns display functional connectivity patterns (Z-transformed correlation, Z(r)) for three representative participants (MSC01–MSC03). A visual inspection revealed consistent topographic distributions across individuals, encompassing canonical large-scale systems, including the default mode, visual, frontoparietal, dorsal attention, language, salience, cingulo-opercular/action-mode, parietal memory, somatomotor, auditory, and somato–cognitive–action networks. Notably, these consistent patterns were observed in both positively and negatively correlated regions, reflecting the canonical architecture of large-scale functional brain systems. The colour scale indicates the functional connectivity strength (Z(r)).

### Convergent and divergent coordination of functional network topography in adult brain

Next, we constructed the functional topography covariance matrix (FOCA matrix) by calculating the correlation efficient between the spatial topography maps of each network pair. Both positive (convergent) and negative (divergent) couplings were retained to systematically delineate the internetwork relationships among all functional networks. The major pattern of the FOCA matrix kept largely unchanged across individuals, characterized by strong intra-network positive couplings (black box) and diversified inter-network couplings (Fig. 2a). We correlated the FOCA matrix between each pair of participants to assess the cross-subject consistency and observed high similarity (mean±std=0.81±0.08; Fig. 2b). Then, we quantified the individual variability of the FOCA matrix (Fig. 2c) by calculating the median absolute deviation (MAD) of the coupling edges across individuals. The variability was relatively low among subnetworks within the DMN, visual and somatomotor networks (mean±std=0.03±0.05) and was markedly high between functional networks, particularly for the internetwork couplings of visual, language and salience networks (mean±std=0.18±0.04; Fig. 2c). Previous studies have reported that stable individualized network mapping largely depends on the total scan time ^[7, 8]^. We therefore evaluated the effect of scan time on FOCA matrix. Iterative split-data analyses demonstrated that the similarity between the matrices of the same individual increased rapidly at very short scan times and gradually reached a plateau (reaching ∼0.80 at 15 min scans and ∼0.85 at 30 min scans; Fig. 2d), and a scan time close to 15 minutes was adequate to derive a reasonable FOCA matrix at the current network resolution.

**Fig. 2.**
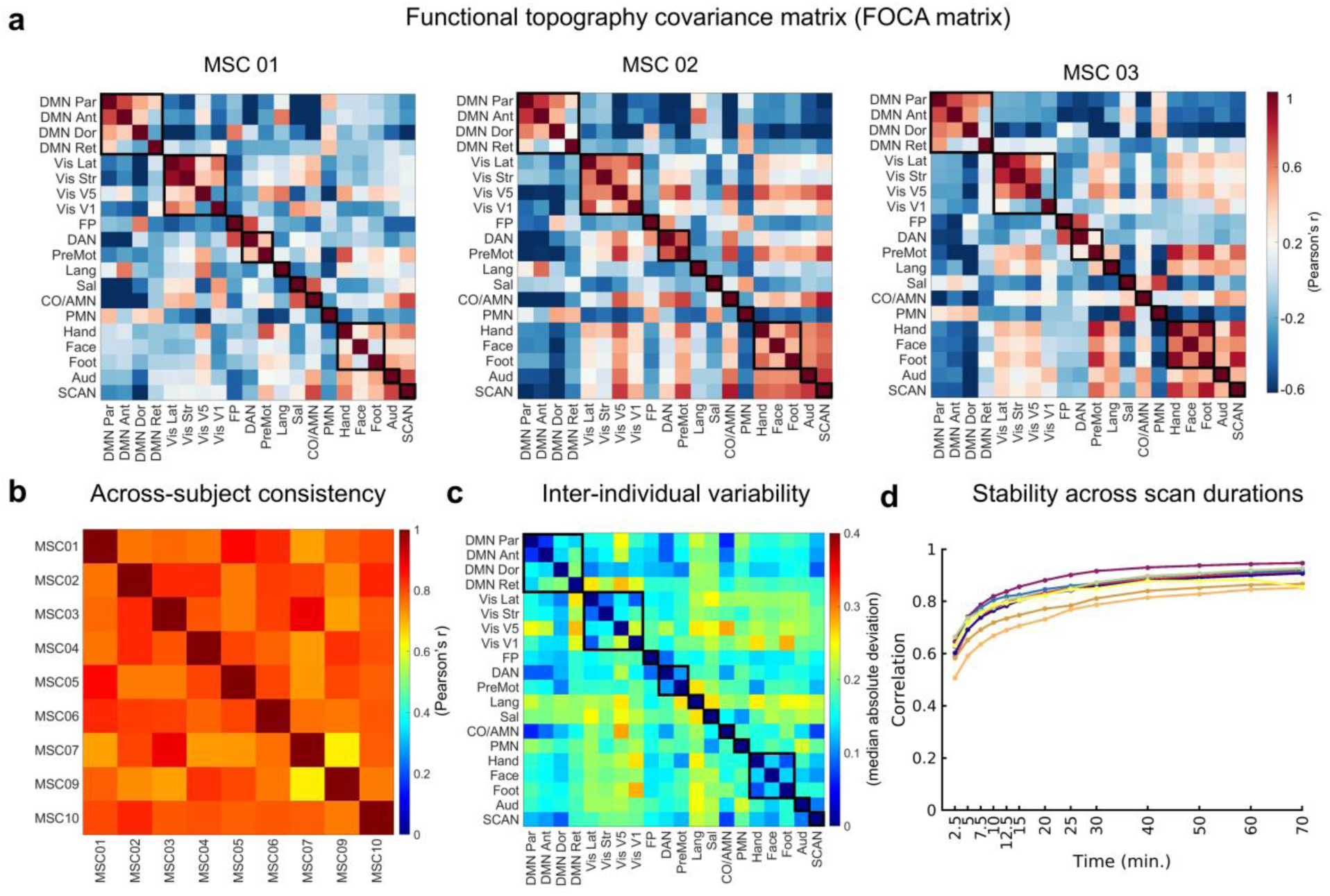
Functional topography covariance matrix (FOCA matrix) for adults. **a,** The individual FOCA matrix for three representative participants (MSC01–MSC03), which captures both positive (convergent) and negative (divergent) couplings. **b,** Across-subject consistency was high, as shown by correlations between the FOCA matrix from different participants. **c,** Interindividual variability, quantified by the median absolute deviation across participants, showed detectable but moderate differences in the FOCA matrix. **d,** Stability across scan durations was evaluated by randomly sampling motion-censored data collected at varying scan durations (x-axis) and comparing it to an independent 70-min dataset from the same subject, which was repeated 1,000 times. Reliability increased rapidly with scan length, with the FOCA matrix similarity reaching ∼0.80 at 15 minutes and ∼0.85 at 30 minutes.

We further applied a hierarchical clustering analysis to delineate the internal organization of the group-level FOCA matrix (for details, see Fig. S6). The feature distance for clustering was defined as one minus the correlation coefficient of topographic maps between each network pair (Fig. 3a). This analysis revealed a prominent hierarchical organization of the FOCA matrix, with six clusters further aggregating into three hyper-clusters (Fig. 3b). Specifically, most clusters were composed of subnetworks from a certain functional system, such as the DMN (Cluster 1: Networks 1–4), visual (Cluster 3: Networks 5–8 and 12), and somatomotor (Cluster 4: Networks 11 and 17–19) systems. Notably, the SCAN clustered with the AMN and auditory network (Cluster 5: Networks 14, 20, and 16) rather than the somatomotor network, which is consistent with its integrative role in linking perceptual and cognitive signals to motor planning ^[1, 35]^. In the second hierarchy, three typical hyper-clusters (namely the internal integrative system, Clusters 1–2; control–attention system, Cluster 6; and primary system, Clusters 3–5) were highly aligned with the previously reported canonical functional gradients from primary to higher-order systems ^[38]^. Notably, higher-order clusters (Clusters 1, 2 and 6; hyper-clusters 1 and 3) exhibited pronounced negative couplings, whereas primary clusters (Clusters 3–5; hyper-cluster 2) exhibited obvious positively couplings (bar plots in Fig. 3b; for each pair of clusters, see Fig. S7). Obvious negative couplings were also identified between higher-order and primary clusters (e.g., between Clusters 1 and 3–5). In figure 3c, we exhibited spatial maps for functional topographies of six representative clusters, along with scatterplots showing intra-cluster (1st row) and inter-cluster (2nd row) couplings. These findings were further validated in the HCP dataset (Figs. S8 and S9).

**Fig. 3.**
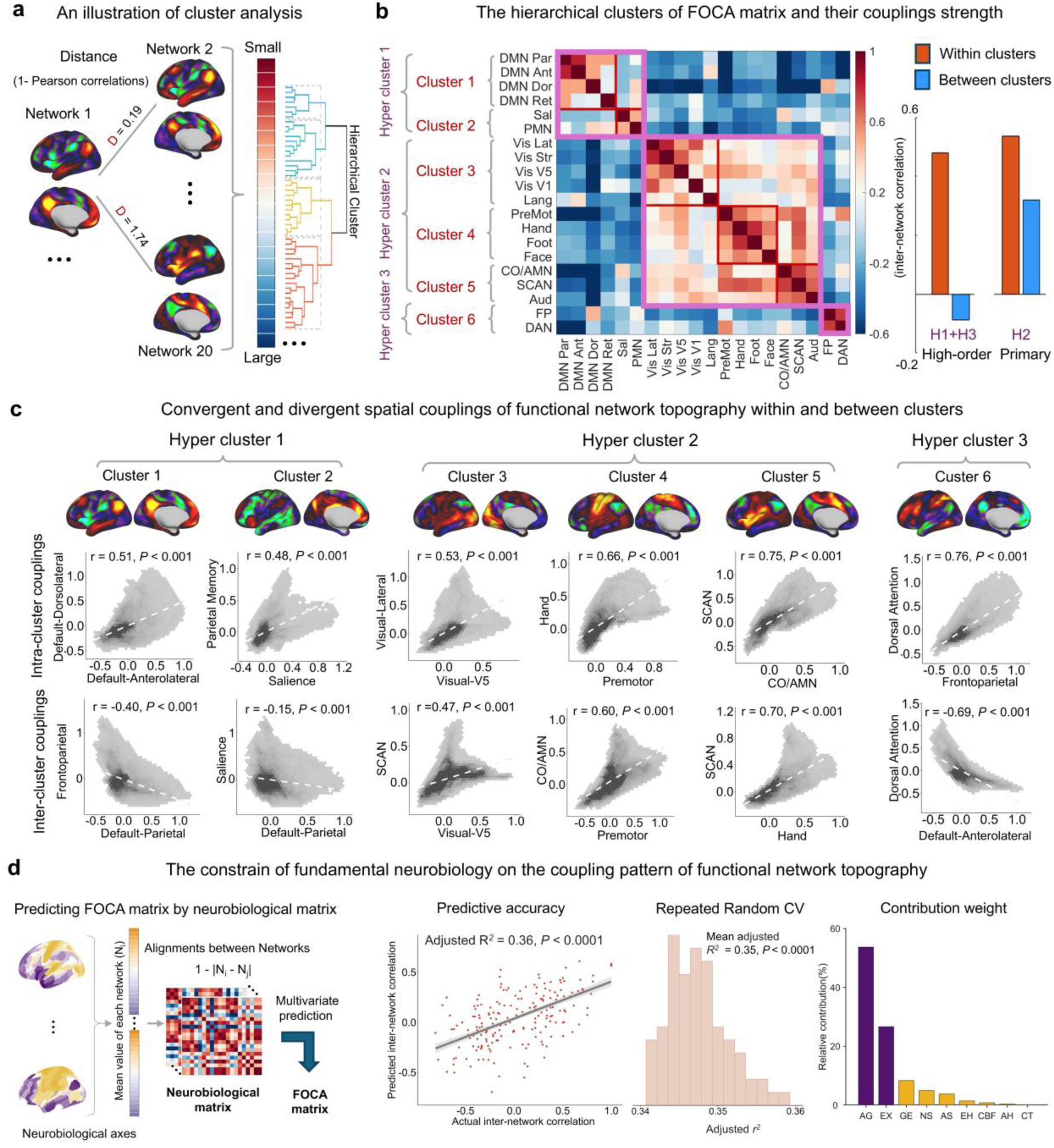
Hierarchical pattern of the topography covariance matrix and its neurobiological basis. **a,** Connectivity profiles in the FOCA group matrix of each network were used as clustering features. **b,** Hierarchical clustering revealed six distinct clusters. Six clusters further formed three hyper-clusters. **c,** Convergent and divergent functional network topographies within and between clusters. The first row shows networks that are positively coupled within clusters, whereas the second row presents networks that are positively coupled between clusters in the primary system (hyper-cluster 2) and negatively coupled between clusters in higher-order systems (hyper-clusters 1 and 3). For each cluster, the top brain maps display the average functional topography of all the constituent networks. **d,** The FOCA matrix is predicted by fundamental brain neurobiological features, with aerobic glycolysis and externopyramidization emerging as the strongest features.

Having established the internetwork relationship between each pair of functional networks, we next examined whether the whole FOCA pattern is constrained by fundamental cortical axes. We employed nine neurobiological maps ^[38]^ representing the anatomical, evolutionary, allometric scaling, aerobic glycolysis, cerebral blood flow, gene expression, neurosynth, externopyramidization, and cortical thickness axes of the brain. We averaged the axis values within each functional network and calculated the alignment (1 minus the pairwise difference) between each network pair to obtain nine neurobiological matrices (Fig. 3d). These matrices were employed as input features of a multivariate regression model to predict the FOCA matrix ^[39]^. We observed a relatively high prediction accuracy, with an adjusted R² = 0.36 (10-fold cross-validation, P < 0.0001), which was robust across 100 repeated random folds (mean adjusted R² = 0.35, P < 0.0001). Further decomposition of feature weights revealed aerobic glycolysis and externopyramidization as the main contributing features (80.3% of the total contribution).

### Distinct spatial coordination of functional network topographies in neonatal brain

Using neonatal scans from the dHCP dataset, we examined whether adult-like FOCA matrix emerge at birth. We re-generated the group-level template (Fig. S10) from a high-quality dHCP subset containing 32 neonates with minimal head motion (mean FD < 0.1). Based on these, we constructed the neonatal FOCA matrix (Fig. 4, Fig. S11) of all individuals. We directly adopted the adult-derived hierarchical clusters into the neonatal matrix as a reference. Compared with adult brain, the neonatal FOCA matrix exhibited a clear hierarchical organization but showed opposite coupling patterns (Fig. 4a). Notably, the visual cluster (Cluster 3) shows strong positive coupling with the DMN cluster (Cluster 1) in neonates rather than with the somatomotor cluster (Cluster 4), as in the adult brain. Higher-order clusters exhibit obvious positive coupling between each other (Clusters 1–2 and 6), in a pattern largely opposite to that in adults (Fig. 4a). At the individual level, the most pronounced developmental differences (Cohen’s d of the difference between the neonatal and adult groups) located in the between-cluster couplings (Fig. 4b) of Cluster 6 (DAN/FP) and Cluster 3 (visual-related networks). Representative values of four network pairs (white asterisks in the left panel; Fig. 4b) are shown in the violin plots (right panel, Fig. 4b). Specifically, the coupling between the DAN and the DMN–Ant as well as the coupling between visual V5 and the DMN-Dor shifted from positive to negative (DAN–DMN: t = 60.72, Cohen’s d = 3.70, *P* < 1×10⁻^12^; visual–DMN: t = 52.89, Cohen’s d = 3.22, *P* < 1×10^⁻12^, the first column). While the coupling between the DAN and AMN as well as the coupling between visual V5 and somatomotor-hand switched from negative to positive (DAN–AMN: t = -51.50, Cohen’s d = -3.14, *P* < 1×10^⁻12^; visual-hand: t = -41.75, Cohen’s d = -2.54, *P* < 1×10^⁻12^, second column). We further visualized their topographic maps and scatterplots of couplings (Fig. 4c, 4d). Notably, we observed evident developmental changes in negative topographies. The location of the most significantly negative connections of the DAN shifted from the primary motor cortex in neonates to the ventromedial prefrontal cortex and precuneus in adults, whereas its positive topography remained largely stable (black arrows, Fig. 4c). The visual network showed similar redistribution of negative connections (black arrows, Fig. 4d) between neonates and adults. We further validated whether these differences were driven by the difference in scan time. By truncating adult scans to the same length of neonates (∼15 min) and repeating the whole analysis, we observed nearly identical results (r = 0.99, *P* < 1×10^-15^, Fig. S12).

**Fig. 4.**
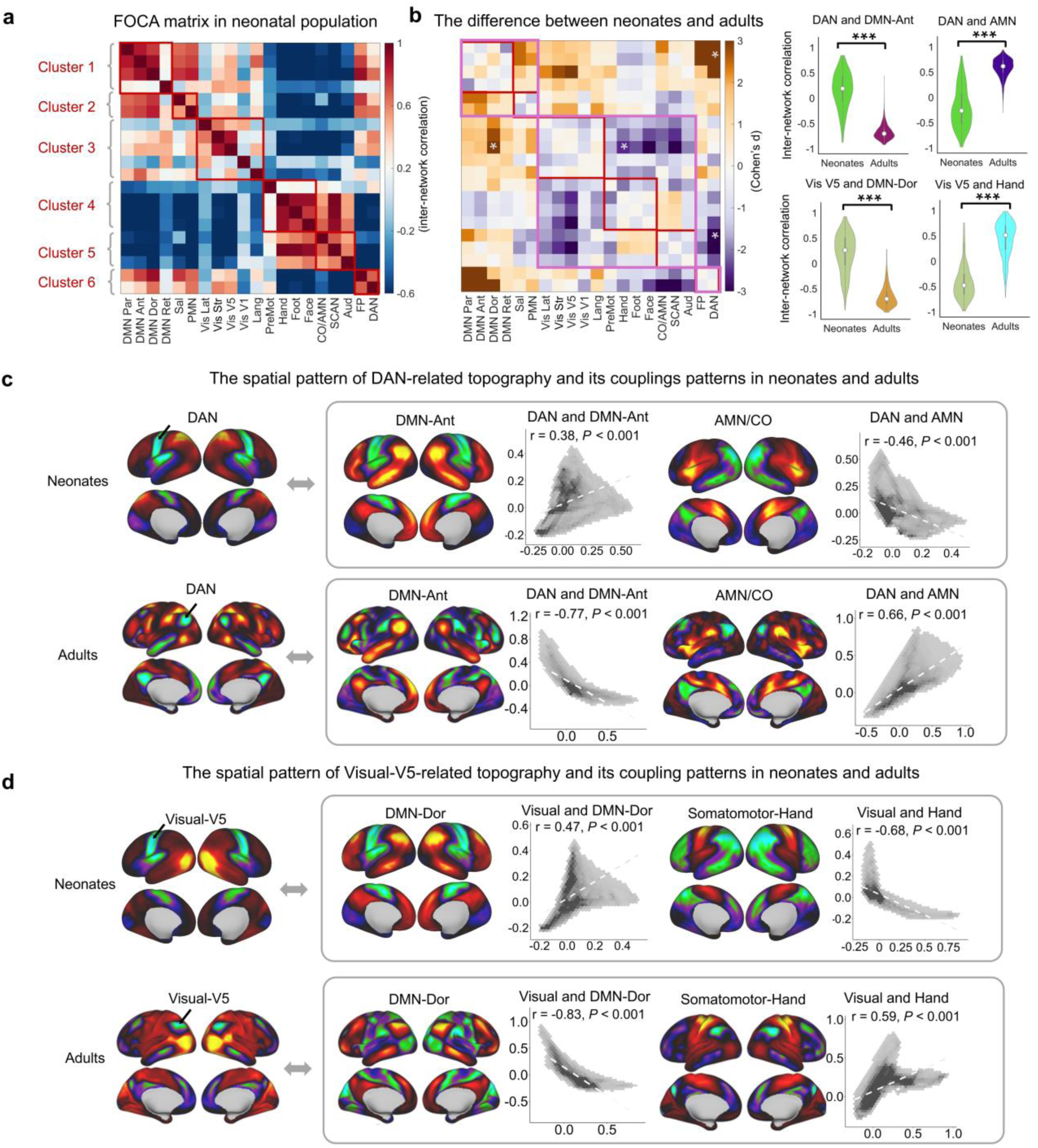
Developmental differences in internetwork covariance between neonates and adults. **a,** FOCA matrix for the neonatal brain showing adult-inverted coupling patterns at the edges between clusters. **b,** Group differences in the FOCA matrices between neonates and adults, with violin plots showing representative changes in four network pairs (white asterisks in the matrix). **c,** The DAN was positively coupled with the DMN but negatively coupled with the AMN in neonates, whereas both relationships were reversed in adults. **d,** The visual V5 network was positively coupled with the DMN but negatively correlated with the somatomotor-hand network in neonates, whereas both relationships were reversed in adults. Notably, these changes were driven primarily by shifts in the negative connectivity (indicated by the black arrows) of the functional topography.

### The spatial alignment of functional topography encodes brain development at birth and predicts neurodevelopmental outcomes at 18 months

Next, we tested whether the person-specific FOCA matrix reflects brain development during the early postnatal period (37 to 44 PMA weeks). We first assessed developmental trajectories in the FOCA matrix using mass univariate generalized additive models (GAMs) controlling for sex, mean framewise displacement (FD), and scan–birth intervals in neonates. The most pronounced developmental effects were observed for inter-cluster couplings (Fig. 5a), especially for those of the auditory network and AMN (white asterisks). Specifically, the couplings of auditory–DMN, auditory–visual, AMN–PMN, and AMN–FP progressively emerged as significantly negative with age, whereas the couplings of auditory–foot and AMN–auditory progressively emerged as significantly positive (Fig. 5b). We further exhibited functional topography maps of two network pairs (auditory–DMN and AMN–FP) at four representative postmenstrual weeks (PMWs) (Fig. 5c). With maturation, the topographies within each network pair became increasingly complementary, accompanied by the redistribution of negative coupling (Fig. 5c). For instance, the angular gyrus gradually exhibited pronounced negative connectivity at 42 weeks for auditory and the AMN network, whereas its positive connectivity already formed at 38 weeks and keeps unchanged (first row for auditory, second row for AMN; Fig. 5c).

**Fig. 5.**
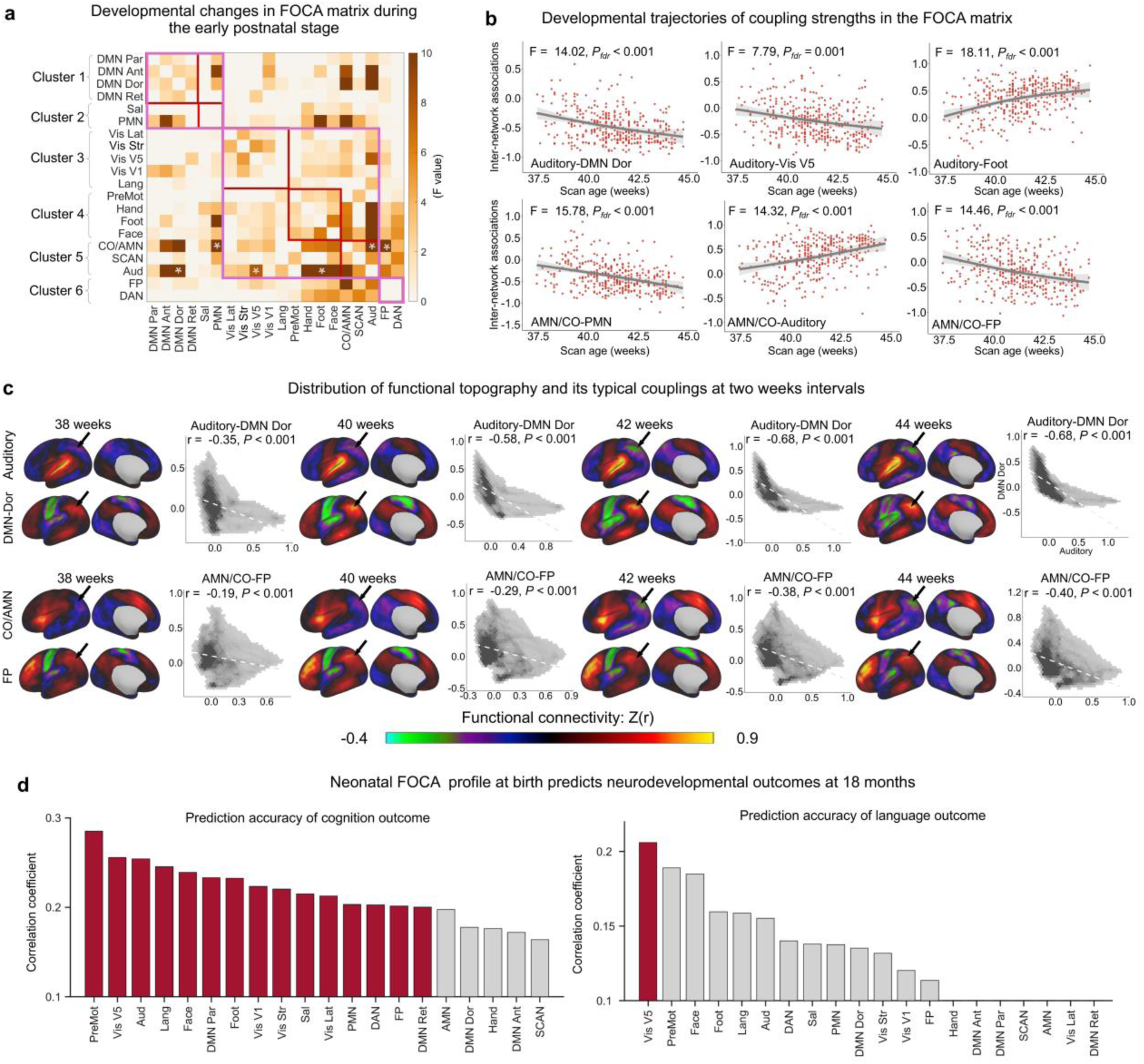
Developmental refinements and neurodevelopmental outcomes of internetwork couplings during the early postnatal period. **a,** Developmental changes in FOCA were assessed using mass univariate generalized additive models (GAMs), controlling for sex, mean FD, and scan–birth intervals. Significant effects were observed across multiple systems, with the auditory network and AMN showing the greatest developmental changes. **b,** Representative developmental trajectories of auditory- and AMN-related couplings. **c,** Representative functional topography maps of auditory- and AMN-related couplings from 37 to 44 PMW. The connectivity pattern progressively becomes complementary to the redistribution of negative couplings in the angular gyrus (indicated by the black arrows). **d,** FOCA profiles at birth predicted neurodevelopmental outcomes at 18 months. Cognitive scores were best predicted by the coupling profiles of premotor and visual-related networks, whereas language scores were significantly better predicted by the coupling profiles of visual V5 networks (FDR-corrected *P* < 0.05). The coloured bars indicate networks with significant predictive accuracy; the grey bars denote nonsignificant results after the FDR correction.

We further examined whether internetwork coordination at birth could predict individual neurodevelopmental outcomes at 18 months. In the dHCP samples, a total of 238 neonates participated in the follow-up assessment, during which cognitive, language, and motor abilities were evaluated using the Bayley Scales of Infant and Toddler Development (Bayley-III, Third Edition). Personalized FOCA profiles from each network were separately entered as features into the multiple linear regression model to predict individual behavioural scores, controlling for sex, mean FD, scan age, and scan–birth intervals (Fig. 5d). We found that 15 of the 20 functional networks could significantly predict cognitive scores at 18 months, with the premotor, visual V5, and auditory networks showing the highest predictive accuracy (FDR-corrected *P* < 0.05). In addition, language scores could also be significantly predicted by the visual V5 network (FDR-corrected *P* < 0.05).

### Comparison between FOCA and conventional FC

A critical issue in the conventional FC analysis is its unstable estimation of negative correlations ^[14, 15]^. FOCA quantified network interactions by assessing the spatial correlations of individualized functional topographies. We conducted three comparison tests to determine whether our approach could outperform conventional FC in representing negative couplings: (1) stability with and without global signal regression (GSR); (2) richness in capturing individual variability; and (3) sensitivity to developmental changes. For conventional FC matrices, we computed connectivity based on the time series of individualized functional networks rather than the group-level network and employed GSR to include a sufficient amount of negative edges to ensure a fair comparison.

First, we computed the mean-normalized RMSE of the matrix between the GSR and non-GSR conditions for each method. Compared with traditional FC, FOCA resulted in a significantly smaller MSE, indicating its stability to GSR (HCP data: t = -25, Cohen’s d = -1.16, *P* < 1×10^-20^; Fig. 6a, left panel). The influences of the two approaches on each pair of networks are also observed in the matrix pattern (Fig. 6a, right panel). Next, we evaluated whether FOCA provides sufficiently differentiated connectivity and more individualized features compared with FC. We assessed the similarity of the negative connectivity profiles for each network between the two methods. We observed substantial dissimilarity, particularly for connections involving the DMN and visual and auditory networks (Table S1; the lowest R² = 0.45). We computed the dissimilarity (1 minus Pearson’s r) between group- and individual-level negative connections to assess individual differences. Compared with FC, FOCA captured significantly greater interindividual variability (t = 22.66, Cohen’s d = 1.03, *P* < 1×10^-20^; Fig. 6b). Finally, we tested whether FOCA captures more developmental effects on negative connectivity than FC. We computed Cohen’s d for the difference between the neonates and adults separately for the FOCA and FC matrices and compared them (FOCA minus FC; Fig. 6c). Compared with FC, the effect size of the FOCA clearly showed an increased difference in development between neonates and adults, particularly for negative couplings involving the DMN and visual and frontoparietal networks (Fig. 6c). Finally, we compared the ability of the FOCA and FC matrices to predict the age of the brain with negative connections. The prediction accuracy was defined as the correlation between predicted brain age (“maturity index”) and chronological age in held-out data (support vector regression, 10-fold cross-validation, controlling for sex, mean FD, and scan–birth intervals). FOCA-based models achieved significantly higher accuracy than FC-based models (FOCA: r = 0.21, *P_perm_* < 0.001; FC: r = 0.11, *P_perm_* = 0.05; Fig. 7d), and this advantage was robust across 100 repetitions of cross-validation (t = 20.38, Cohen’s d = 2.88, *P* < 1×10^-20^; Fig. 7d). Together, these findings establish FOCA as a robust and development-sensitive framework for characterizing internetwork relationships that significantly exceeds conventional FC.

**Fig. 6.**
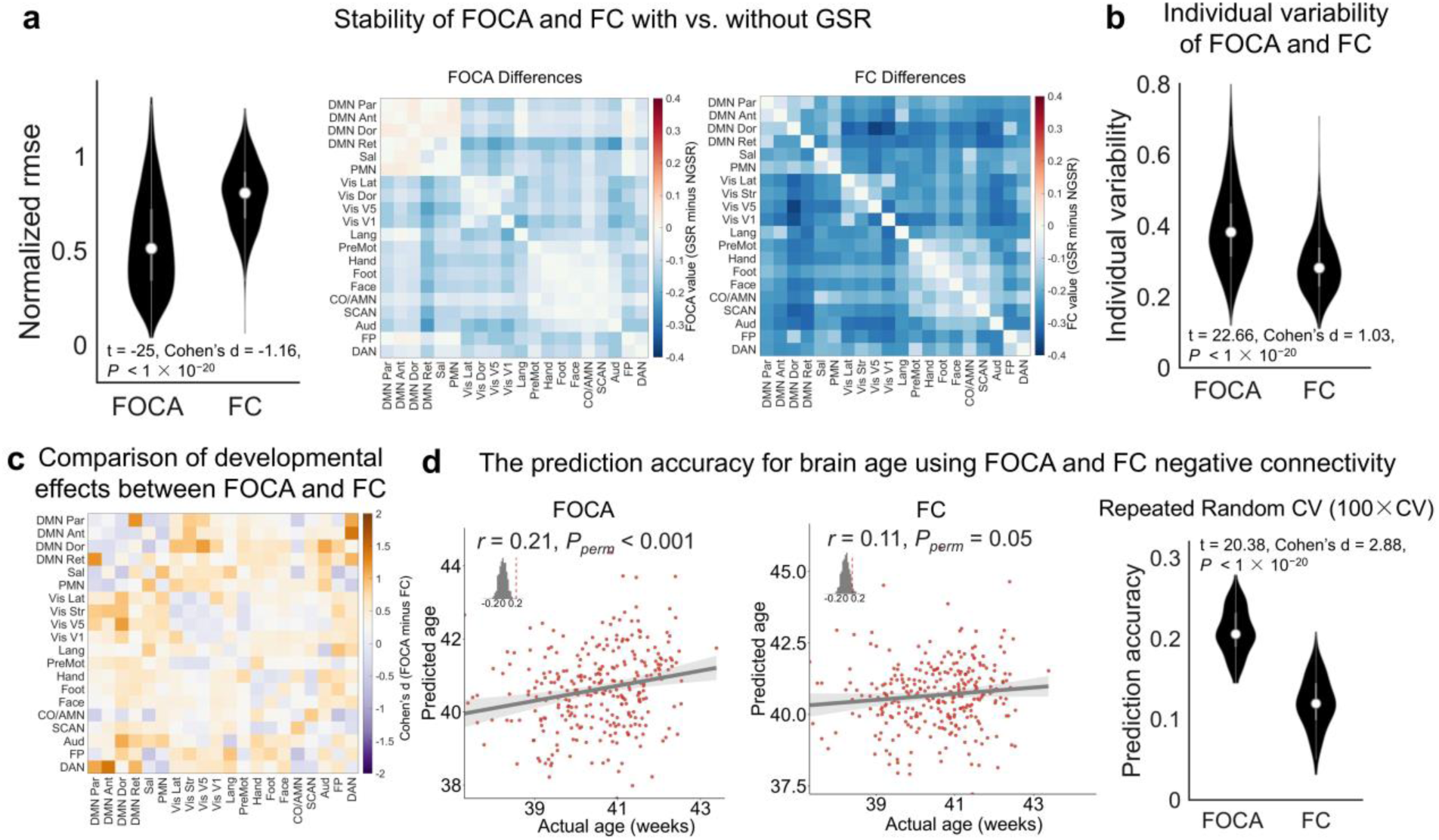
FOCA provides unique and more stable information beyond conventional FC. **a,** Stability of FOCA and FC with and without GSR. The stability of FOCA was significantly greater than that of FC, as shown by the lower normalized RMSE between the GSR and non-GSR conditions. Heatmaps show differences in FOCA and FC induced by GSR. **b,** Individual variability in FOCA and FC. Violin plots show interindividual dissimilarity (1 minus Pearson’s r) between group- and individual-level negative connections. FOCA captured significantly stronger individual variability than FC. **c,** Differences in the developmental effects captured by FOCA versus FC. Compared with FC, FOCA could capture substantially greater developmental effects, particularly in negative couplings involving the DMN and visual and frontoparietal networks. **d,** Prediction of early brain maturation. Scatter plots show correlations between predicted and chronological ages using FOCA (left panel) or FC (middle panel) features. FOCA achieved higher accuracy than FC. The violin plot (right panel) shows that this advantage was robust across 100 repetitions of cross-validation.

### Sensitivity analysis

We assessed the stability of our findings across three major dimensions. (i) cross-dataset replication: individualized topographies and FOCA matrices derived from an independent adult cohort (HCP) closely matched those from MSC at both group and individual levels, indicating high reproducibility of the hierarchical organization (Fig. S9). (ii) global signal regression: FOCA was highly consistent with and without GSR in adults and neonates, for both topographies and inter-cluster coupling profiles, indicating its high robustness (Fig. S13, S14). (iii) scan-duration harmonization: truncating adult HCP scans into neonatal-length showed little influence on the neonate-adult differences of FOCA, supporting that our results were not driven by the short scan time in neonates (Fig. S12). These validations converge to show that FOCA’s internetwork architecture and its developmental inferences are relatively reliable to methodological variations.

## Discussion

We proposed FOCA as a novel framework to systematically quantify the internetwork relationships between individualized functional topographies and their developmental origins. We identified a reproducible, self-clustered hierarchical architecture of brain FOCA matrix. Typical coupling patterns exist among functional systems and are well predicted by the fundamental neurobiological axis. Neonatal networks exhibit adult-inverted coupling patterns among functional hierarchies and significant adjustments of coupling profiles in auditory and action-mode networks, driven by obvious spatial redistributions of negative connectivity. Neonatal FOCA profiles, especially in primary visual networks, could significantly predict individual neurodevelopmental outcomes at 18 months. Moreover, in terms of capturing negative couplings, FOCA exhibits advantages in the robustness to global signal regression and the sensitivity to age-related maturation compared with conventional FC. These findings identify biologically meaningful internetwork alignments of the individualized functional network along the cortex and its developmental remodelling at the neonatal stage, establishing a novel and reliable analytical framework for probing the convergent and divergent relationships between brain functional systems.

Prior work on large-scale networks has emphasized spatial organization and functional roles across the lifespan ^[25, 40]^, but network interactions—especially negative correlations—have received relatively little attention. The human brain is intrinsically organized into large-scale systems that often exhibit cooperative and antagonistic activity patterns ^[1, 2]^, yet traditional definitions based on time series correlations are highly sensitive to preprocessing choices, especially GSR, which mathematically enforces anticorrelations and has fuelled long-standing controversy ^[15]^. Although some researchers view these anticorrelations as artefacts, others have provided compelling evidence for their neurobiological relevance ^[41, 42]^; for example, antagonistic coupling between the dorsolateral prefrontal cortex and subgenual anterior cingulate predicts a clinical improvement in depression ^[42]^. Recent studies have further demonstrated that anticorrelated activity observed during diverse tasks is similar to that observed at rest ^[41]^ and that patterns of negative connectivity differ markedly between infants and adults ^[27, 43]^. We introduce a topography-based approach that quantifies network interactions through spatial patterns of functional network, rather than raw time series correlations to resolve this debate. FOCA provides a robust, development-sensitive, and predictive characterization of internetwork organization—especially for negative interactions—that conventional FC fails to capture. This framework reveals that the anticorrelated architecture is a prominent axis of developmental change and demonstrates a robust antagonistic structure independent of GSR.

Our findings reveal pronounced developmental differences in the FOCA matrix between neonates and adults, with adult brains exhibiting robust negative correlations among higher-order networks (e.g., the DMN and DAN) as well as between higher-order networks and primary networks (e.g., the DMN and visual network; the DMN and somatomotor network), whereas neonates display predominantly positive relationships. This divergence likely reflects the protracted maturation of substrates that enable functional competition: long-range, heavily myelinated frontoparietal/frontotemporal pathways supporting rapid state switching ^[44]^; the integrity of thalamocortical control loops; and a finely tuned excitatory–inhibitory balance, particularly GABAergic inhibition, that permits cross-network suppression ^[45, 46]^. In neonates, these elements are immature (limited myelination, underdeveloped long-range connectivity, incomplete thalamic regulation, and weaker inhibition), and early behaviour favours global integration across sensory, motor, and associative systems rather than modular specialization ^[47, 48]^. Consistent with this finding, visual and language networks are negatively coupled with somatomotor/action systems in neonates but positively coupled in adults, paralleling the emergence of coordinated visuomotor behaviour and speech–motor control. Similarly, PMN–language coupling shifts from positive in neonates to negative in adults, indicating a transition from broad coupling to functional specialization that reduces interference and improves the efficiency of language processing and contextual memory retrieval. Together, these reversals indicate that the emergence of robust internetwork competition is a hallmark of later network specialization and the maturation of cognitive control.

A key mechanistic insight from our analyses is that aerobic glycolysis is the strongest predictor of the FOCA matrix. Aerobic glycolysis is most pronounced in association cortices that act as integrative hubs ^[49]^, supporting long-range connectivity and higher-order cognitive functions. Unlike primary sensory and motor areas, which are metabolically more efficient and developmentally precocious, association cortices rely on aerobic glycolysis to sustain elevated synaptic remodelling, plasticity, and persistent activity ^[50]^. This metabolic specialization likely provides the energetic substrate required for cross-network coordination. Consistent with this view, increased aerobic glycolysis is associated with evolutionary cortical expansion ^[51]^ and prolonged postnatal maturation, features aligned with the developmental refinement and flexibility of the human association cortex ^[52]^. Together, these observations suggest that the disproportionate contribution of aerobic glycolysis reflects not only energetic demands but also a neurobiological mechanism enabling integrative and adaptive large-scale network interactions.

In our developmental trajectory analyses, we observed that the auditory network and the action-mode network exhibited the most pronounced changes in topographic associations with other networks across the neonatal period. This pattern likely reflects both the maturational timetable and the functional role of these systems. The auditory network undergoes relatively early functional specialization, shaped by extensive prenatal and postnatal exposure to sound, and progressively integrates with higher-order cognitive networks ^[53]^. In contrast, the action-mode network—encompassing the anterior cingulate, insular, and prefrontal regions—serves as a high-order control hub, orchestrating goal-directed actions and coordinating interactions with sensory and motor systems ^[1]^. Its topographic coupling is therefore highly sensitive to structural and functional remodelling during early life. Biologically, both networks include regions with rapid myelination, a high metabolic demand, and substantial aerobic glycolysis during this stage ^[54]^, which may accelerate their reconfiguration and integration with other large-scale systems. The prominence of these two networks in our results suggests that early sensory processing and action-related control functions constitute major axes of neonatal brain network reorganization.

Our framework of functional network topographic mapping opens several promising avenues for future research. First, it provides a foundation for charting the developmental trajectory of large-scale network interactions at different stages across the human lifespan ^[55]^. Second, by stably capturing the spatial alignments of network topographies, our approach may increase the sensitivity and reproducibility in linking brain functional architecture, especially antagonistic functional patterns to behavioural phenotypes ^[56, 57]^. Finally, the whole mapping of internetwork relationships derived from functional topography may provide new evidence and insights into refined, individualized connectome targets, especially functional inhibitory circuits for neuromodulation, potentially increasing the precision of interventions for affective disorders ^[34, 58]^.

Despite these strengths, several limitations should be acknowledged. First, although large-scale topographic mapping provides a robust framework for characterizing the individualized brain organization, a finer-grained delineation of functional networks would provide a richer description of internetwork relationships than a network resolution of 20. However, such high-resolution individualization remains technically challenging given the current constraints on the scan duration, signal-to-noise ratio, and computational resources. Second, the observed inversion of functional hierarchy in neonates raises intriguing mechanistic questions. Understanding why early functional organization exhibits this opposite hierarchical pattern will require dedicated modelling efforts and integrative approaches combining computational simulations, developmental neurobiology, and longitudinal imaging data. Third, the scan duration of fMRI in neonates inherently shorter than that in adults. Previous studies have reported that achieving stable individualized functional parcels in neonates may require long fMRI scans due to greater noise and motion ^[27]^. Interestingly, our validation analyses showed that FOCA matrices exhibited a convergent spatial pattern within only 15 minutes of scanning. These findings suggest that compared with vertex-level maps, certain derived indices of individualized functional parcellation, such as spatial covariance matrices at a large scale, may have lower scan demands. Moreover, our FOCA matrix effectively captured individual differences related to brain maturation and later behavioural outcomes, indicating that meaningful interindividual variability can be detected at the current scan time. Nevertheless, characterizing the stability of FOCA matrices in long fMRI acquisitions in infants is necessary and important in the future. Together, our findings provide a new framework for understanding key fundamental principles underlying the emergence, antagonism and maturation of the functional architecture of the human brain.

## Materials and methods

### Participants and data acquisition

The MSC dataset included 10 individuals (5 females, aged 24–34 years), with 300 minutes and 12 resting-state fMRI sessions, as previously described ^[7]^. Participant MSC08 was excluded from this study due to high levels of head motion and self-reported sleep, leaving nine participants in final analysis. All participants provided written informed consent. The procedures were approved by the Washington University Institutional Review Board and School of Medicine Human Studies Committee.

The HCP S1200 dataset comprised 996 adults (532 females, aged 22–37 years) ^[31]^, with 58-minute resting-state fMRI. Written informed consent was obtained from all participants, and the scanning protocol was approved by the Institutional Review Board of Washington University in St. Louis, MO, USA (IRB #20120436).

The dHCP ^[30]^ included high spatial-temporal resolution data from 370 term-born neonates (173 females; gestational age [GA] at birth = 37–42 weeks; postmenstrual age [PMA] at scan = 37–44 weeks; 15 minutes scan of resting-state fMRI). All infants were scanned during natural sleep without sedation to obtain structural and resting-state fMRI data using a 3T Philips Achieva scanner. Written consent was obtained from the guardian of each baby.

### MRI data preprocessing

The MSC dataset was obtained online (https://openneuro.org/) in a preprocessed, fully denoised and surface-registered format, and no further preprocessing or denoising was performed in the present study.

All functional images in the HCP dataset initially underwent minimal preprocessing using the HCP public pipeline ^[59]^. The final stage of preprocessing was performed using GRETNA v2.0.0, including smoothing, linear detrending, nuisance variable regression, scrubbing, and temporal filtering. Notably, for Friston’s 24 head motion parameters, average signals were removed using multivariate linear regression, and spike regression-based scrubbing (0.5 mm displacement) was performed to control for the effect of head motion. Finally, the denoised fMRI time series was spatially smoothed using Gaussian kernels with a full width at half maximum (FWHM) of 6 mm (σ = 2.55 mm), applied geodesically for surface data via the Connectome Workbench command-line utilities.

For all the fMRI images in the dHCP dataset, the minimal preprocessing steps were performed by the dHCP organizer and have been validated in prior studies ^[60]^. Building upon this framework, we extended the preprocessing steps to the cortical surface by projecting each participant’s rs-fMRI time series from the native volumetric space to the native cortical surface using a ribbon-constrained volume-to-surface mapping approach. Surface-projected time series were subsequently resampled to the 40-week PMA symmetric surface template via MSM-based native-to-template registration. Importantly, due to the relatively coarse spatial resolution of neonatal fMRI data (2.15 mm isotropic), which exceeds the typical neonatal cortical thickness (∼1 mm), substantial partial volume effects are present within grey matter (GM) voxels. Therefore, the standard HCP pipeline is not directly applicable to dHCP data. The dHCP pipeline incorporates a volumetric partial volume correction (PVC) procedure (see https://git.fmrib.ox.ac.uk/seanf/dhcp-neonatal-fmri-pipeline/-/blob/master/doc/surface.md?ref_type=heads) that adjusts for GM contamination in the functional signal prior to surface mapping to address this issue. Following surface projection and registration, postprocessing steps identical to those used in the HCP dataset were applied, including spatial smoothing, nuisance signal regression, scrubbing, and temporal filtering. Spatial smoothing was performed using Gaussian kernels with a full width at half maximum (FWHM) of 6 mm (σ = 2.55 mm).

### Generating individualized functional networks based on the TM method

We first computed individualized functional networks using a widely applied template-matching (TM) approach to determine the functional network topography ^[27, 32, 33]^. The reference atlas comprises 20 group-level functional networks defined in previous studies ^[7, 34–37]^ (Fig. S2), reflecting the current consensus on large-scale brain organization. These networks included the default mode (DMN, Networks 1–4), visual (Networks 5–8), frontoparietal (Network 9), dorsal attention (DAN, Networks 10–11), language (Network 12), salience (Network 13), cingulo-opercular/action-mode (CO/AMN, Network 14), parietal memory (PMN, Network 15), auditory (Network 16), somatomotor (Networks 17–19), and somato–cognitive–action networks ^[35]^ (SCAN, Network 20).

The group-level functional network served as a reference for defining networks and generating seed-based correlation maps for each participant. For each network, the seed time series was defined as the mean time series within the group-level network mask. Individual seed-based correlation maps were averaged across participants in the MSC dataset and thresholded at z > 1 (corresponding to the top ∼15.9% of connections), yielding the MSC-derived templates (Fig. S3). These templates were subsequently applied to generate individualized functional networks for participants in both the MSC and the HCP datasets. Whole-brain functional connectivity matrices were computed for each grayordinate; as an illustrative example, we visualized the connectivity profile of a single grayordinate (Fig. S4). For each grayordinate, connectivity maps were thresholded using the same criteria as the group template, and η² values were computed to quantify the proportion of variance in the connectivity profile explained by each network. Each grayordinate was then assigned to the network with the highest η² value.

For the neonatal dataset (dHCP), a group-level functional template was first constructed from a high-quality subset of 32 neonates with minimal head motion (mean FD < 0.1 mm) to ensure robustness (Fig. S10). Individualized functional networks were then derived for each neonate using the same TM procedure, ensuring methodological consistency across developmental cohorts.

### Convergent and divergent functional network topography in adults

We first derived participant-specific topographic maps by correlating the seed time series of each individualized functional network (mean of that in all network vertices, Pearson’s r, Fisher z–transformed) to characterize interactions among large-scale functional networks. This procedure yields a connectivity map (including both positive and negative correlations) for each network that reflects its spatial topography. Based on these individualized functional topography maps, we constructed a FOCA matrix by computing Pearson’s correlation coefficients between the maps of all network pairs. Positive correlations were interpreted as convergent couplings, whereas negative correlations reflected divergent couplings.

Across-subject consistency was assessed by computing pairwise correlations of the FOCA matrix between participants. Interindividual variability was quantified using the median absolute deviation (MAD), a robust nonparametric measure of dispersion ^[10]^. Iterative split-data analyses ^[7, 8]^ were performed to examine the effect of the scan acquisition length on reliability. For each subject, motion-censored data segments of varying lengths were randomly selected and split into independent halves, where the FOCA matrix was separately estimated. The similarity between the two halves was quantified using Pearson’s correlation analysis and repeated 1,000 times per period.

We averaged the functional topographic maps across participants to generate group-level representations for each network and to summarize convergent and divergent topographic patterns. We computed pairwise Pearson’s correlation coefficients between all network-specific functional maps and constructed a FOCA matrix to quantify the interactions among networks. Hierarchical clustering was then applied to the FOCA matrix to identify large-scale patterns of network interactions. Clustering was performed using the average distance method and distance metric (1 minus Pearson’s correlation coefficient), implemented in MATLAB R2020b. Inter-cluster relationships were further summarized by averaging the intra- and inter-cluster correlations, enabling the assessment of large-scale modular organization.

The same analytic pipeline was applied to an independent cohort from the HCP dataset to evaluate replicability. Group- and individual-level measures of the FOCA matrix were compared across datasets to assess generalizability. Finally, we repeated the analysis using the HCP dataset with and without GSR and compared the resulting functional topographies and FOCA matrices to examine the robustness to preprocessing choices.

### Internetwork covariance constrained by fundamental organizational features of the brain

We incorporated nine cortical neurobiological axes ^[38]^, including the anatomical hierarchy, quantified by the T1w/T2w ratio; evolutionary hierarchy, quantified by macaque-to-human cortical expansion; allometric scaling, quantified as the relative extent of areal scaling with the overall brain size; aerobic glycolysis, quantified from PET measures of oxygen consumption and glucose use; cerebral blood flow, quantified via arterial spin labelling; gene expression, quantified by the first principal component of genes expressed in the brain; neurosynth, quantified by the first principal component of meta-analytic decoding; externopyramidization, quantified as the ratio of supragranular to infragranular pyramidal soma size; and cortical thickness, quantified from structural MRI (HCP S1200 dataset), to determine whether internetwork covariation is shaped by the fundamental organizational features of the cortex.

For each brain map, network-by-network similarity matrices were derived by averaging feature values within each network and calculating the feature alignment matrix by estimating the alignment (1 minus the pairwise difference) between each pair of networks (Fig. 3d). These matrices were subsequently entered into a multiple linear regression model to explain the observed network interactions of the functional network topography. Model performance was evaluated using 10-fold cross-validation, with the adjusted R^2^ serving as the primary metric of prediction accuracy. The procedure was repeated across 100 random folds to assess robustness. Finally, the relative feature weights of each neurobiological axis were computed to determine their contributions to explaining the FOCA matrix.

### Convergent and divergent functional network topography in neonates

Using the individualized functional networks for each neonate, we generated corresponding functional topographic maps. For each neonate, the FOCA matrix was computed by correlating the spatial distributions of all network pairs. We applied the adult-derived hierarchical clustering solution to the neonatal matrices and assessed the correspondence of the FOCA matrix across individuals to enable direct comparisons throughout development. Group- and individual-level analyses were performed to assess the correspondence of the FOCA matrix during development. At the individual level, the similarity to group-level reference matrices was quantified by calculating Pearson’s correlation coefficients.

Finally, we examined the robustness of the neonatal results to global signal regression by recomputing the FOCA matrix using data processed with and without GSR and assessed the similarity of the resulting matrices.

### Developmental refinement and neurodevelopmental outcomes of internetwork covariance during the early postnatal period

We investigated the developmental changes in internetwork covariance during the early postnatal period by modelling age-related trajectories of the FOCA matrix in neonates using mass univariate GAMs. For each network pair among the 20 functional networks, correlation coefficients were fitted as a smooth function of PMA at scan, while controlling for sex, mean FD, and scan–birth intervals. Significant age-related effects were identified by examining the fitted GAM coefficients across network pairs, with multiple comparisons corrected using the false discovery rate (FDR). We characterized age-specific patterns by constructing group-level functional topographies for each PMA week between 37 and 44 weeks. This approach enabled the visualization of week-by-week changes in the convergent and divergent functional topographies captured by the FOCA matrix.

We tested whether neonatal internetwork covariance would predict later outcomes by following 238 neonates who underwent a behavioural assessment at 18 months using the Bayley Scales of Infant and Toddler Development, Third Edition (Bayley-III). FOCA profiles from each network at birth were entered as predictors in multiple linear regression models, with cognitive, language, and motor scores as dependent variables. All the models controlled for sex, mean FD, scan age, and scan–birth intervals. The statistical significance of the regression coefficients was assessed using the FDR correction (*q* < 0.05).

### Comparisons between FOCA and conventional FC

We compared FOCA against conventional FC in three aspects: (i) stability to preprocessing choices, (ii) shared vs. unique information content, (iii) developmental sensitivity, and predictive validity. First, we calculated the mean-normalized root mean square error (RMSE) between connectivity matrices generated with and without GSR, normalized by the mean value of the matrix, to compare the stability of FOCA with that of conventional FC. Group-level differences in stability were assessed using two-sample t tests. Then, we computed the coefficient of determination (R²) between the FOCA and FC profiles and focused on negative connections to evaluate whether FOCA provides information beyond FC. In addition, subject-level dissimilarity was quantified as 1 minus Pearson’s r between group- and individual-level negative connections, and differences between methods were assessed using paired t tests. Next, the developmental effect was assessed by comparing FOCA- or FC-derived connectivity between neonates and adults using two-sample t tests, with effect sizes estimated using Cohen’s d. Last, to confirm the predictive validity of FOCA or FC features, linear support vector regression (SVR) models were trained to predict chronological age from FOCA or FC features during the neonatal period. Prior to prediction, confounding variables (sex, mean FD, and scan-to-birth interval) were regressed out from both the features and chronological ages. The prediction accuracy was defined as the Pearson correlation coefficient between the predicted and actual age in held-out folds. Robustness was confirmed by repeating the 10-fold cross-validation 100 times, and group-level differences in predictive performance were assessed using paired t tests.

## Data availability

All data required for reproducing our findings have been publicly available, including the individualized functional networks, Group- and individual-level functional topography, FOCA matrix, and the data for visualizing main figures. They are stored in a publicly accessible cloud repository (github: https://github.com/zhaohuaxishi1/Convergent_Divergent_FOCA_Development). The MSC data are publicly available at https://openneuro.org/datasets/ds000224. For the HCP dataset, raw image scans are publicly available at https://www.humanconnectome.org/. Source data are provided with this paper. For the dHCP dataset, raw image scans are publicly available at https://nda.nih.gov/.

## Code availability

Software packages used in this manuscript include HCP pipeline (https://github.com/Washington-University/HCPpipelines/releases/), dHCP structural pipeline (https://github.com/BioMedIA/dhcp-structural-pipeline), dHCP functional pipeline (https://git.fmrib.ox.ac.uk/seanf/dhcp-neonatal-fmri-pipeline/-/tree/master), Connectome Workbench (https://www.humanconnectome.org/software/connectome-workbench), cifti-matlab toolbox v2 (https://github.com/Washington-University/cifti-matlab/), SPM12 toolbox (https://www.fil.ion.ucl.ac.uk/spm/software/spm12/), GRETNA toolbox v2.0.0 (https://www.nitrc.org/projects/gretna/), Template matching v 1.0 (https://github.com/DCAN-Labs/compare_matrices_to_assign_networks), MSM (https://github.com/ecr05/MSM_HOCR/releases), LIBSVM (3.25) (https://www.csie.ntu.edu.tw/~cjlin/libsvm/), Support Vector Regression (https://github.com/ZaixuCui/Pattern_Regression_Clean), and R 4.0.3 (https://www.r-project.org), Matlab 2020b (https://www.mathworks.com/products/matlab.html). The codes used in this study are available at github: https://github.com/zhaohuaxishi1/Convergent_Divergent_FOCA_Development.

## Acknowledgments

This work was supported by the National Natural Science Foundation of China (Nos. 82327807), and the Fundamental Research Funds for the Central Universities (Nos. 2233300002, 2233100018).

## Author Contributions

J.L.Z., Y. Z and T.D.Z. designed research; T.D.Z., and J.L.Z designed the methodology. J.L.Z. developed visualizations. L.L.S, and Y.H.X provide methodological guidance; J.L.Z. performed the data analysis; J.L.Z., and T.D.Z. wrote the paper; J.L.Z. and T.D.Z. revised the paper.

