## Supplementary Information for "Convergent and divergent spatial topographies of individualized brain functional networks and their developmental origins"

### Supplementary figures

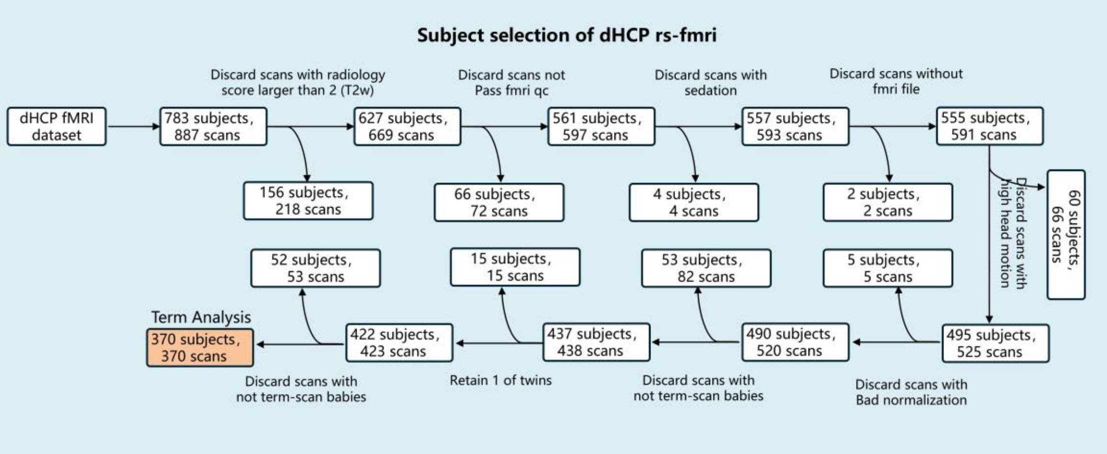


**Fig. S1 Flowchart illustrating inclusion/exclusion criteria.** Of the initial 783 participants, 156 were excluded based on low radiological scores (ranging from 3 to 5). 66 participants were excluded due to failure to pass the fMRI quality control. 4 participants and 2 participants were excluded because of sedation and lack of fMRI files, respectively. 60 participants were excluded due to excessive head motion (head-motion displacement >5 mm, rotation >5°, or mean frame wise displacement (mFD) > 0.5 mm). 5 participants were excluded due to poor registration quality. 53 participants were excluded because they were born prematurely and not term-scan. 15 participants were excluded due to being twins. 52 participants were excluded due to preterm-born neonates at term-equivalent age. After rigorous quality control, a total of 370 participants remained.

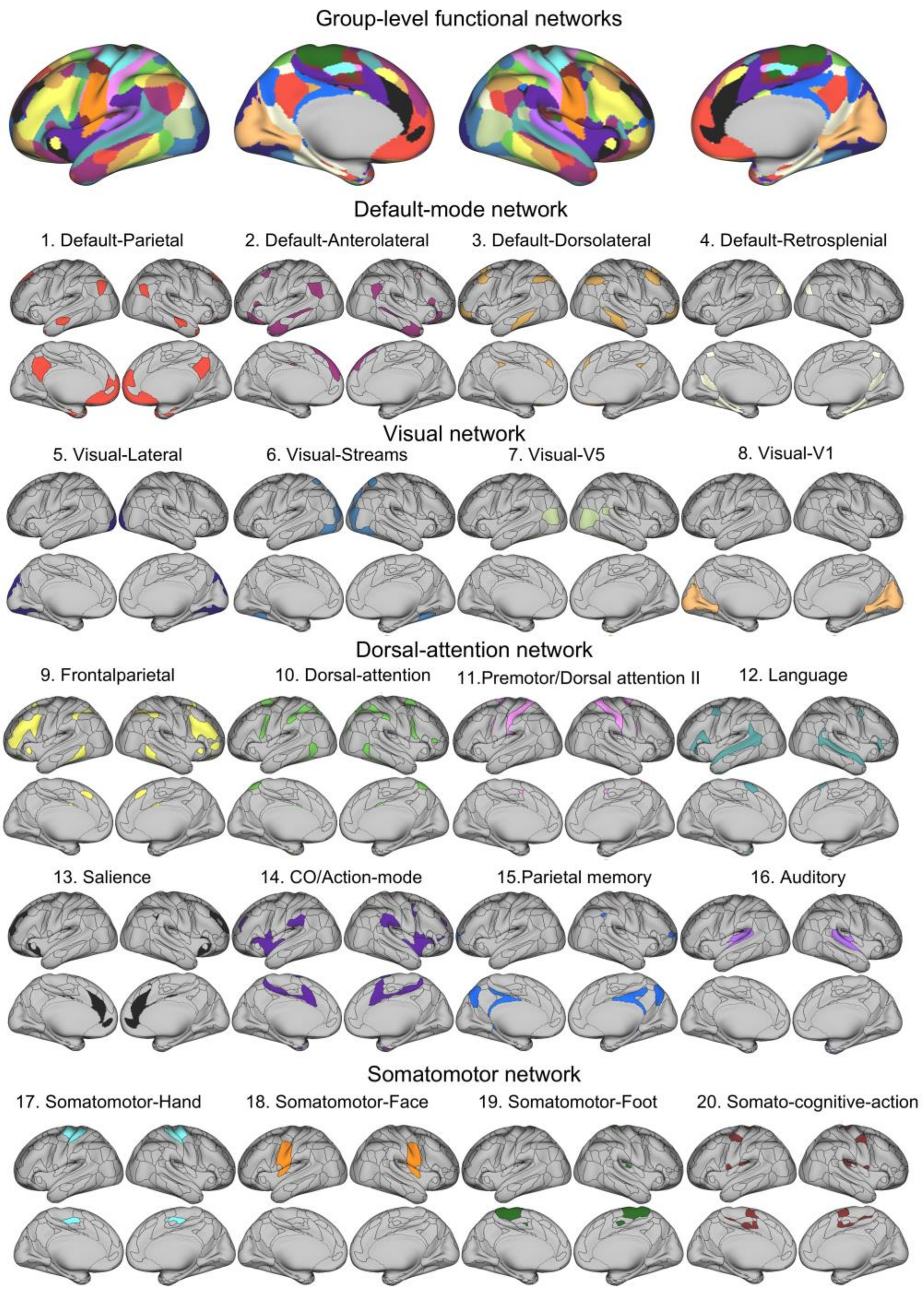


**Fig. S2 Group-level functional networks**. we adopted a recently proposed fine-grained atlas comprising 11 major functional networks with 20 subnetworks[1-5]. These networks include the default mode network (DMN, Networks 1–4: parietal, anterolateral, dorsolateral, and retrosplenial), the visual network (Networks 5–8: lateral, streams, V5, and V1), the frontoparietal network (Network 9), the dorsal attention network (Networks 10-11), the language network (Network 12), the salience network (Network 13), the cingulo-opercular/action mode network (CO/AMN, Network 14), the parietal memory network (Network 15), the auditory network (Network 16), the somatomotor network (Networks 17–19: hand, face, and foot), and the somato-cognitive-action network (SCAN, Network 20).

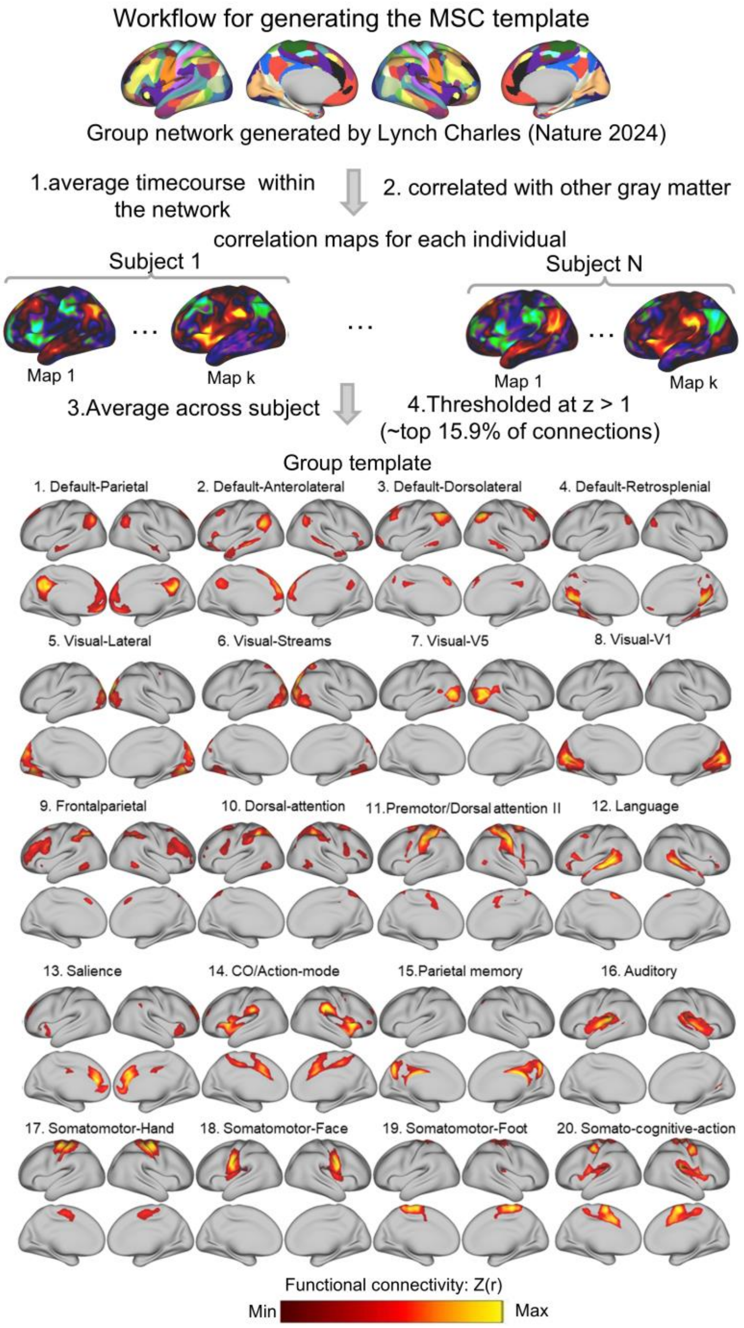


**Fig. S3 Generation of MSC-based group templates using seed-based correlation mapping.** Group-level functional networks were used as references to generate group templates. For each network, the average time course within the network was extracted (Step 1) and correlated with all other grayordinates to generate whole-brain seed-based correlation maps for each individual (Step 2). These individual correlation maps were then averaged across participants in the MSC dataset to create group-level correlation templates for each network. Templates were thresholded at $z > 1$ (~top 15.9% of connections) to define the final group-level functional template for networks such as the default mode network (DMN), cingulo-opercular/action-mode (CO/AMN), and somatomotor–hand (Hand SM).

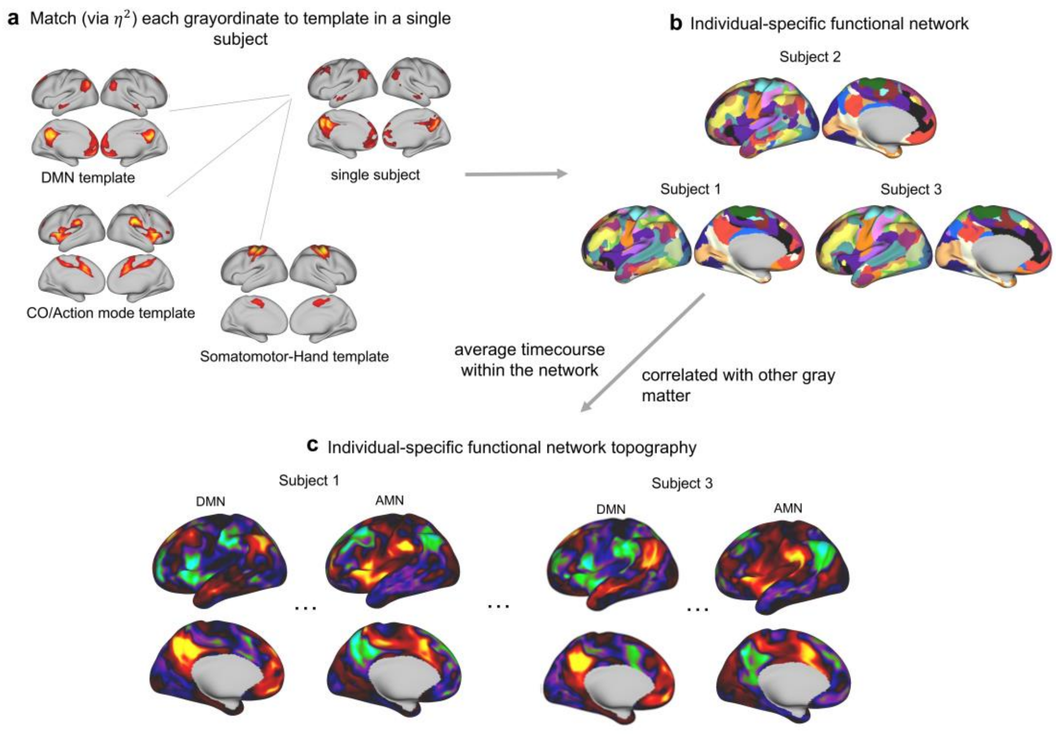


**Fig. S4 Pipeline for generating individualized functional networks and functional topography using the template-matching (TM) approach. a,** For each participant, each grayordinate's connectivity profile was compared to MSC-derived network templates, and $\eta^2$ values were computed to quantify similarity. Each grayordinate was assigned to the network with the maximum $\eta^2$ value, resulting in individualized parcellations. **b,** Individual-specific network maps were generated for all participants. **c,** To derive functional network topography, the mean time series within each individualized network was extracted and correlated voxel-wise with the entire brain, producing participant-specific maps that characterize spatial distribution of each network.

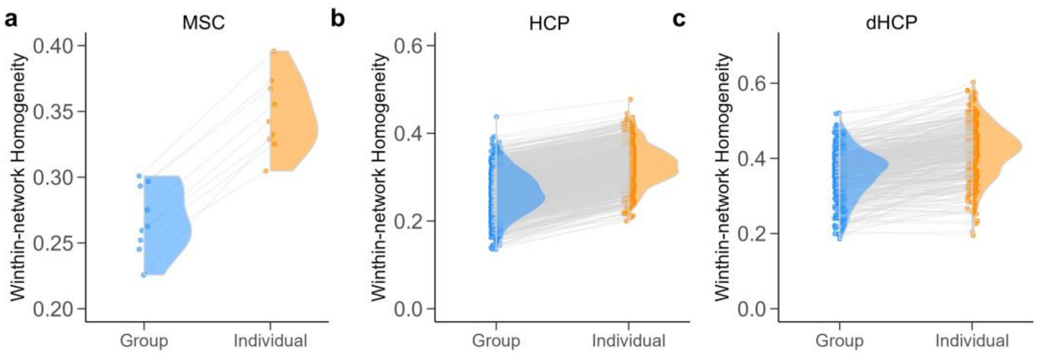


**Fig. S5 Individualized networks exhibited significantly higher within-network homogeneity compared to group-level networks across all datasets. a,** In the MSC dataset, individualized functional networks showed markedly greater within-network homogeneity compared with group-level networks (t = 6.23, $P = 1.20 \times 10^{-5,}$ Cohen's d = 5.69). **b,** Similar improvements in functional coherence were observed in the adult HCP dataset (t = 31.49, $P < 1.0 \times 10^{-5}$, Cohen's d = 4.48). **c,** The neonatal dHCP dataset demonstrated a consistent pattern, with individualized parcellations yielding higher within-network homogeneity than group-level references (t = 12.56, $P < 1.0 \times 10^{-5}$, Cohen's d = 1.45). Each gray line represents a paired comparison for a single participant.

.

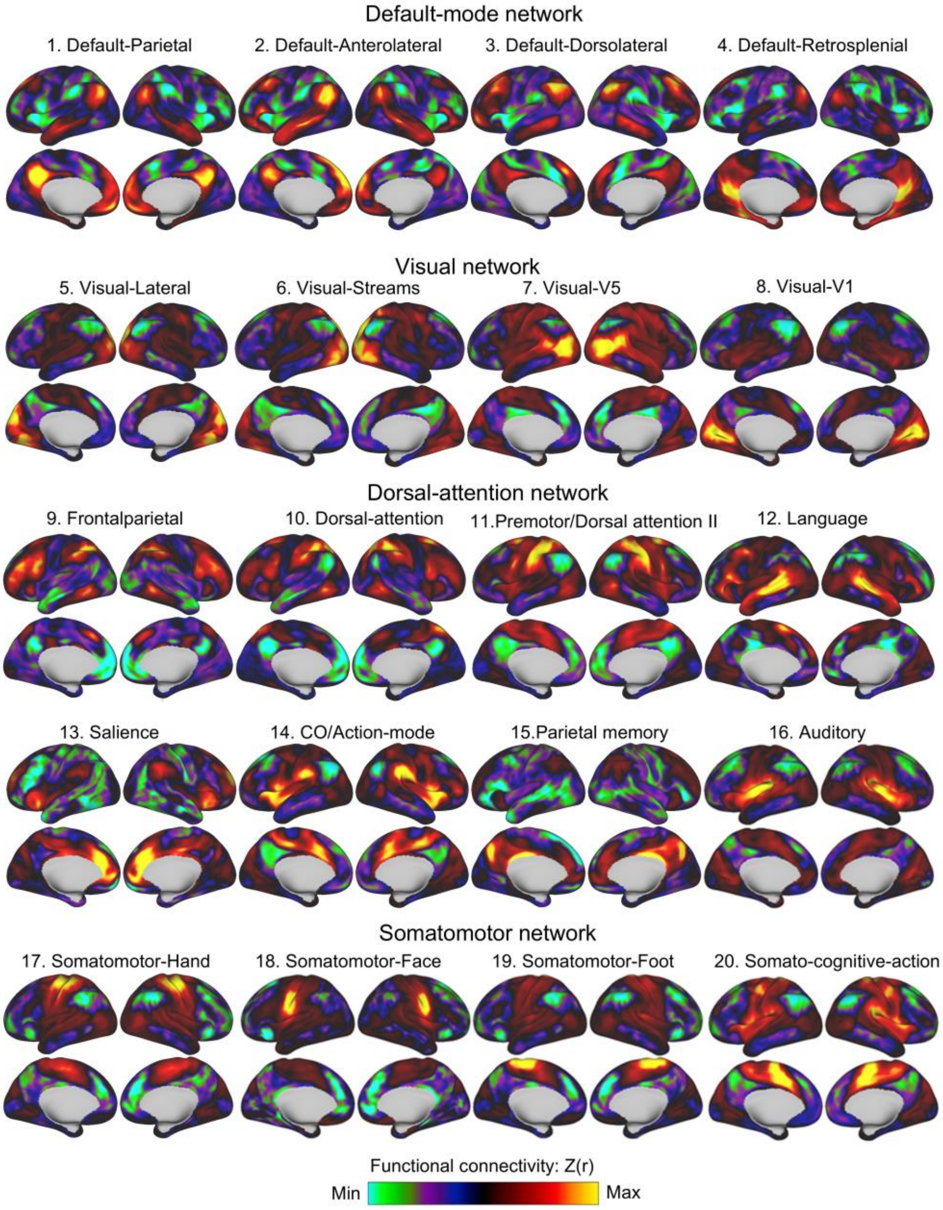


**Fig. S6 MSC-based group-level probabilistic representations of 20 large-scale functional networks.** Group-level probabilistic maps were generated by averaging topographic representations across participants in the MSC dataset. These networks reflect the current consensus on large-scale functional brain organization and include the default mode network (DMN, Networks 1–4: parietal, anterolateral, dorsolateral, and retrosplenial), the visual network (Networks 5–8: lateral, streams, V5, and V1), the frontoparietal network (Network 9), the dorsal attention network (Networks 10–11), the language network (Network 12), the salience network (Network 13), the cingulo-opercular/action mode network (CO/AMN, Network 14), the parietal memory network (Network 15), the auditory network (Network 16), the somatomotor network (Networks 17–19: hand, face, and foot), and the somato-cognitive-action network (SCAN, Network 20). Color scale indicates functional connectivity strength within each network (green = minimum, red = maximum).

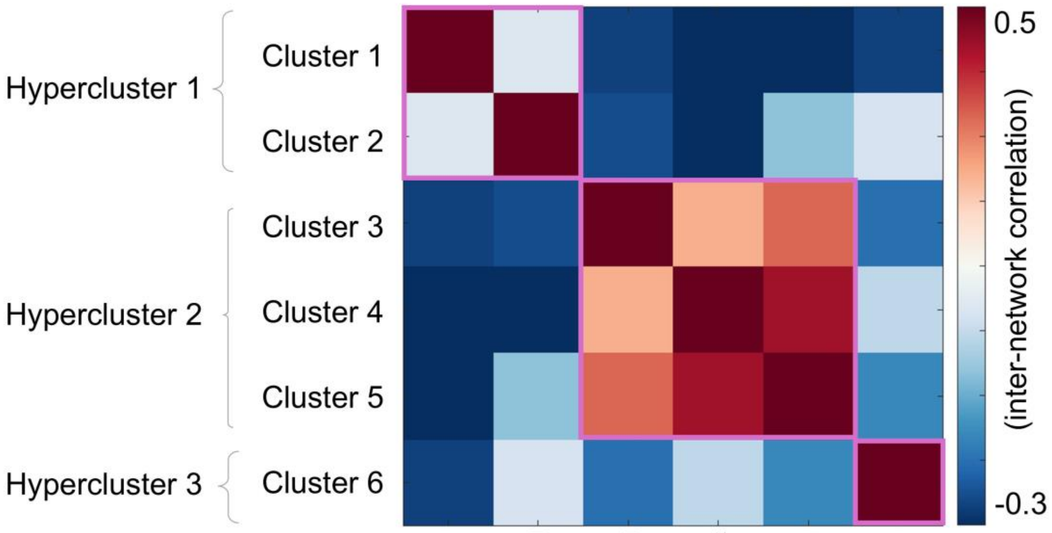


**Fig. S7 Hyper-clusters organization of inter-cluster relationships.**
By averaging intra- and inter-cluster correlations, we derived intercluster connectivity patterns, which revealed a hierarchical hyper-cluster structure. Specifically, Cluster 1 and Cluster 2 grouped together as hypercluster 1, Clusters 3–5 formed hypercluster 2, and Cluster 6 constituted hypercluster 3, indicating that fine-grained clusters can be further integrated into three broader modules.

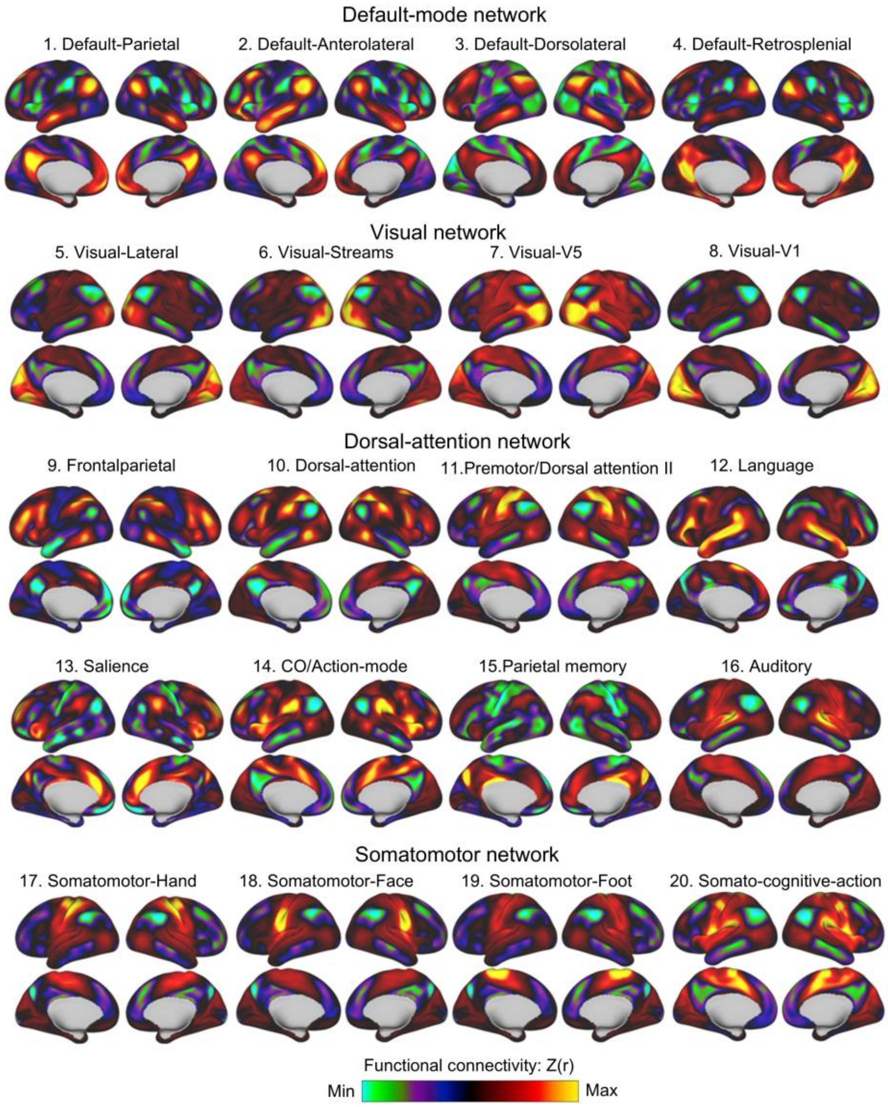


**Fig. S8 HCP-based group-level probabilistic representations of 20 large-scale functional networks.** Group-level probabilistic maps were generated by averaging topographic representations across participants in the HCP dataset. These networks reflect the current consensus on large-scale functional brain organization and include: the default mode network (DMN, Networks 1–4: parietal, anterolateral, dorsolateral, and retrosplenial), the visual network (Networks 5–8: lateral, streams, V5, and V1), the frontoparietal network (Network 9), the dorsal attention network (Networks 10–11), the language network (Network 12), the salience network (Network 13), the cingulo-opercular/action mode network (CO/AMN, Network 14), the parietal memory network (Network 15), the auditory network (Network 16), the somatomotor network (Networks 17–19: hand, face, and foot), and the somato-cognitive-action network (SCAN, Network 20). Color scale indicates functional connectivity strength within each network (green = minimum, red = maximum).

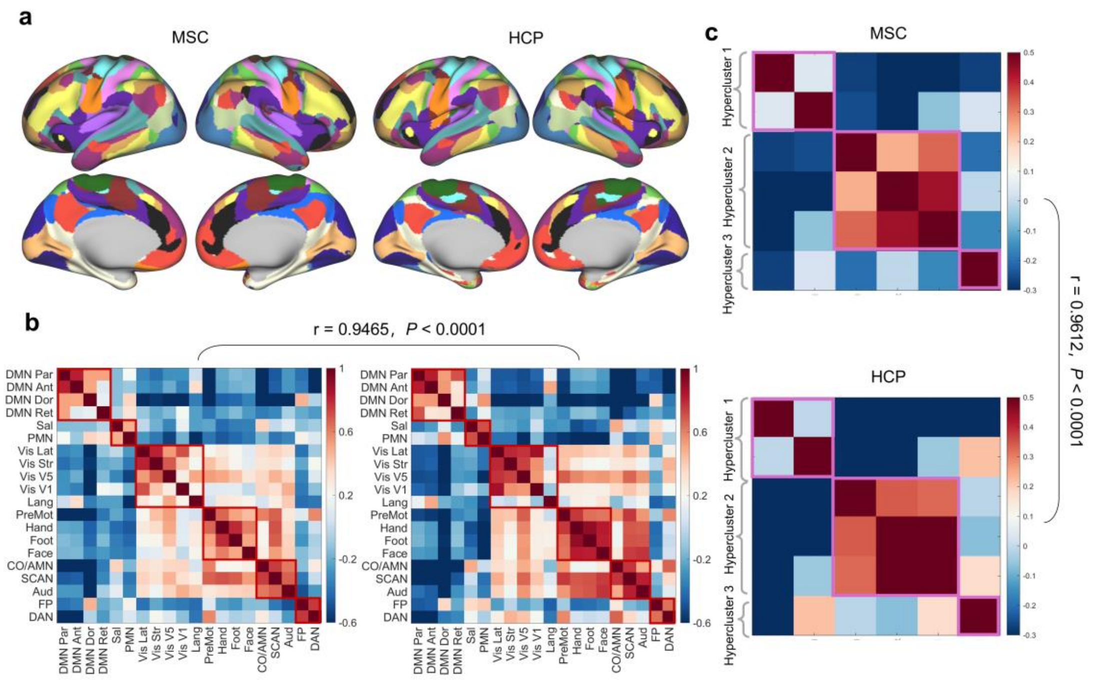


**Fig. S9 Stability of functional network topography across independent adult cohorts. a,** Group-level probabilistic topographies were converted into discrete network assignments by labeling each cortical vertex according to its highest network loading. Shown are group-level functional network parcellations for the MSC dataset (left) and HCP dataset (right), both derived using the same individualized template-matching procedure. FOCA matrices were obtained by averaging topographic maps across participants within each cohort. **b,** FOCA matrix was highly similar across cohorts (r = 0.9465, $P < 0.0001$). **c,** Inter-cluster correlation matrices was highly similar across cohorts (r = 0.9612, $P < 0.0001$), demonstrating strong reproducibility of large-scale topography organization in adults. Warm colors indicate positive coupling , and cool colors indicate negative coupling between functional network topography.

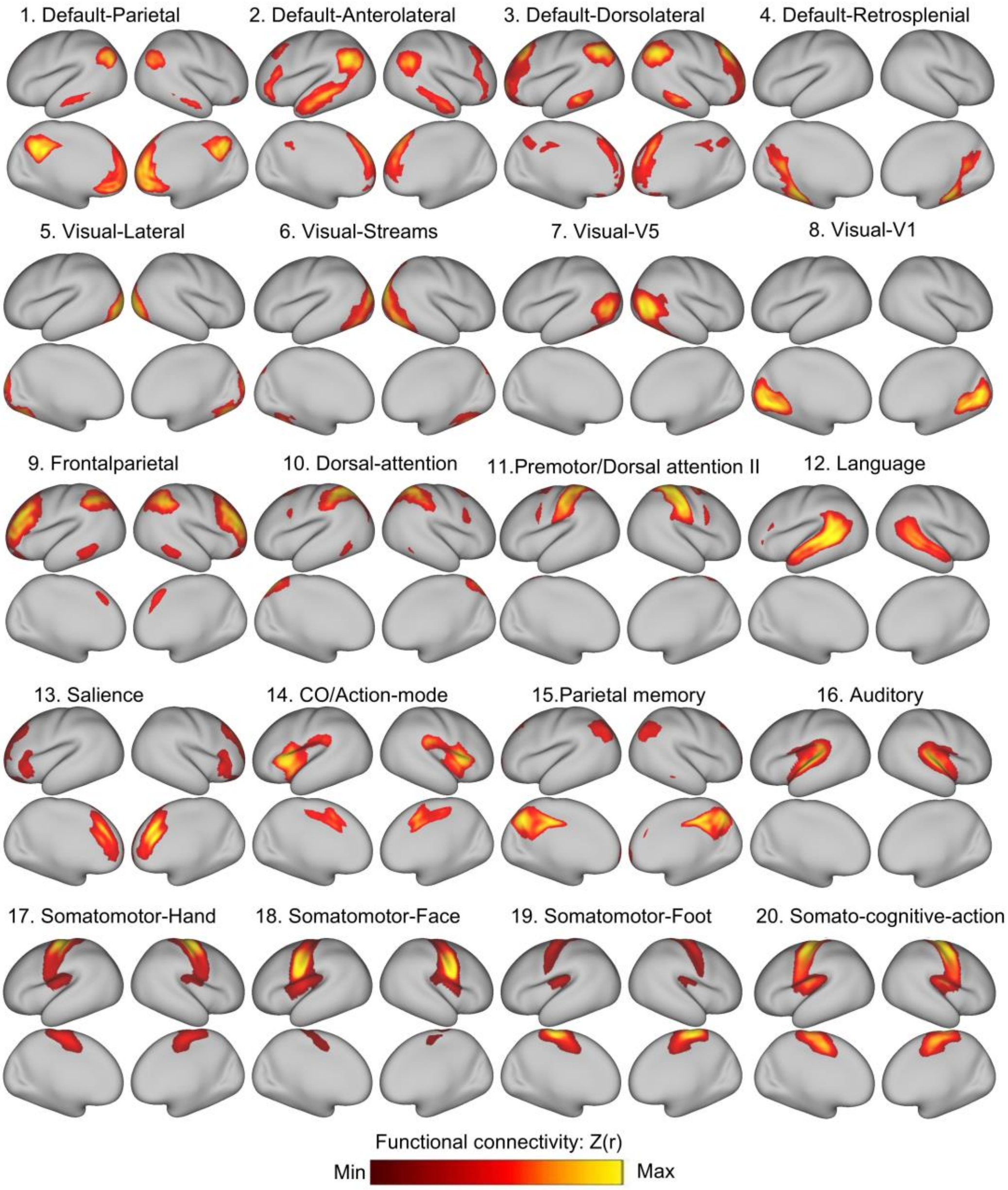


**Fig. S10 Group-level template of dHCP dataset.** A group-level template was first generated from a high-quality subset of 32 neonates with minimal motion (mean FD < 0.1). These networks template encompass the default mode network (DMN, Networks 1–4: parietal, anterolateral, dorsolateral, and retrosplenial), the visual network (Networks 5–8: lateral, streams, V5, and V1), the frontoparietal network (Network 9), the dorsal attention network (Networks 10–11), the language network (Network 12), the salience network (Network 13), the cingulo-opercular/action mode network (CO/AMN, Network 14), the parietal memory network (Network 15), the auditory network (Network 16), the somatomotor network (Networks 17–19: hand, face, and foot), and the somato-cognitive-action network (SCAN, Network 20).

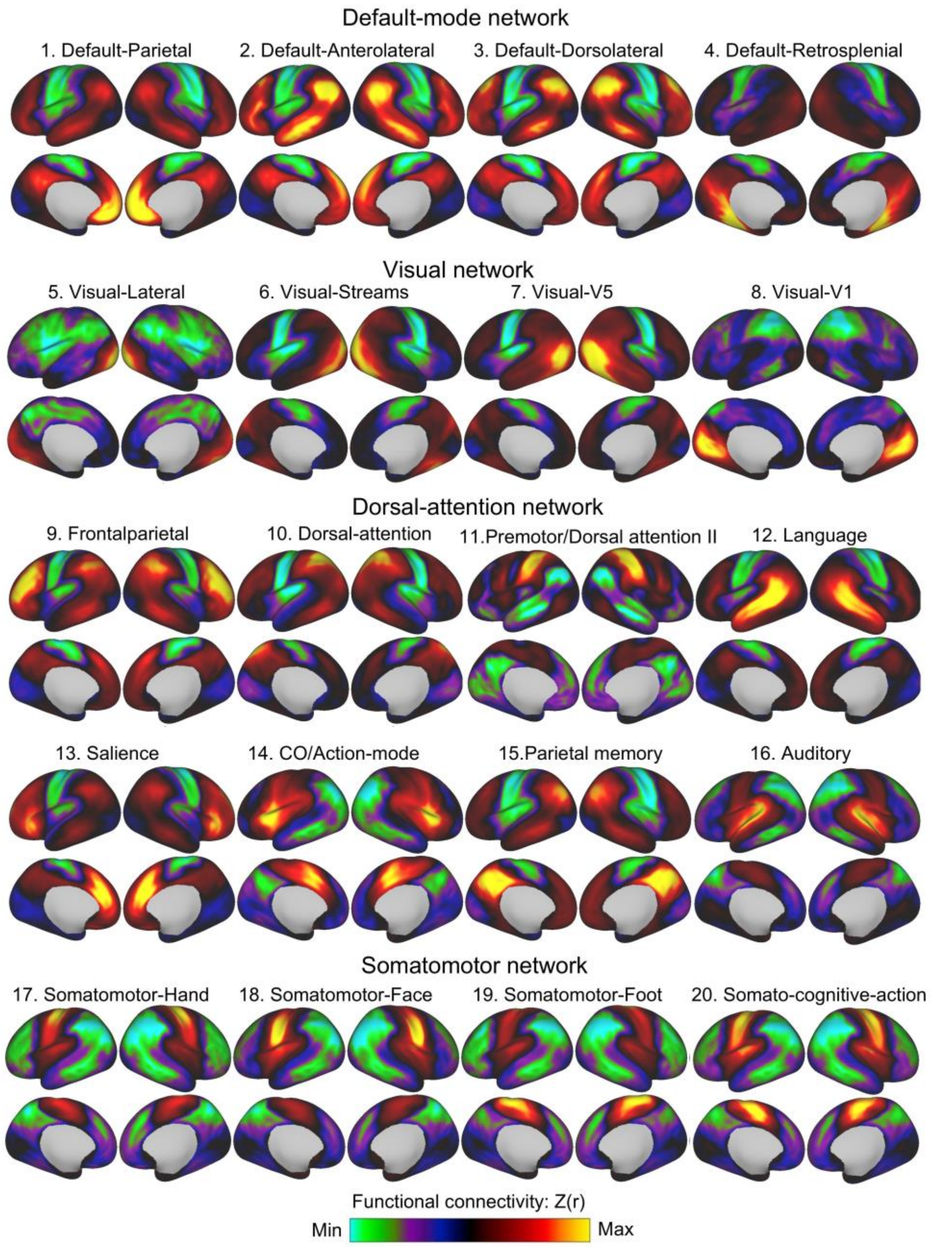


**Fig. S11 dHCP-based group-level probabilistic representations of 20 large-scale functional networks.** Group-level probabilistic maps were generated by averaging functional topographic representations across participants in the dHCP dataset. These networks include: the default mode network (DMN, Networks 1–4: parietal, anterolateral, dorsolateral, and retrosplenial), the visual network (Networks 5–8: lateral, streams, V5, and V1), the frontoparietal network (Network 9), the dorsal attention network (Networks 10–11), the language network (Network 12), the salience network (Network 13), the cingulo-opercular/action mode network (CO/AMN, Network 14), the parietal memory network (Network 15), the auditory network (Network 16), the somatomotor network (Networks 17–19: hand, face, and foot), and the somato-cognitive-action network (SCAN, Network 20).

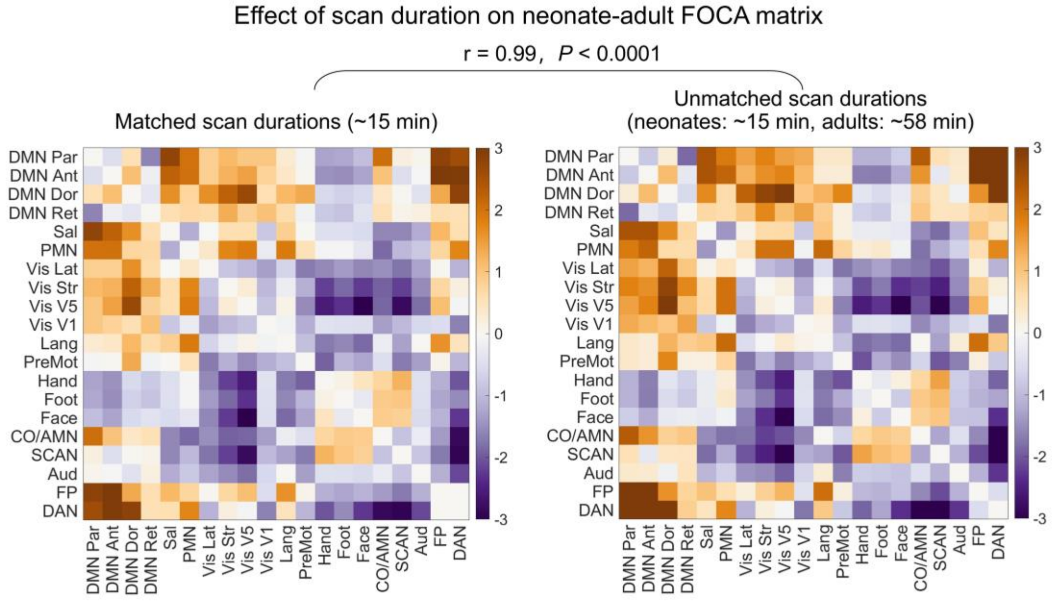


**Fig. S12 Effect of scan duration on neonate-adult differences in FOCA matrix.** FOCA matrices were compared between neonates and adults using either matched (~15 min) or unmatched (neonates: ~15 min; adults: ~58 min) scan durations. The spatial patterns of group differences were nearly identical between the two approaches ($r = 0.99$, $P < 0.0001$), indicating that developmental differences in FOCA matrix are not driven by scan length.

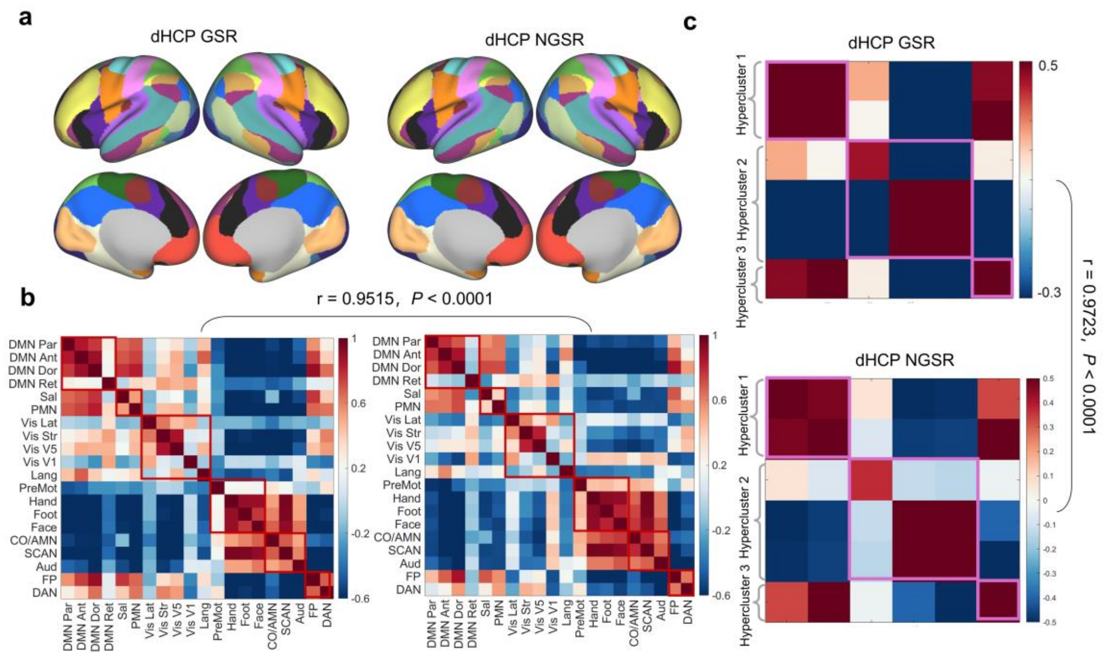


**Fig. S13 Robustness of neonatal functional network topography to global signal regression (GSR). a,** Group-level probabilistic topographies were converted into discrete network assignments by labeling each cortical vertex according to its highest network loading. Shown are group-level functional network parcellations for the dHCP dataset computed with GSR (left) and without GSR (NGSR, right). **b,** The FOCA matrices were highly similar between the GSR and non-GSR conditions. **c,** The intercluster correlation matrices also showed high similarity with and without GSR, demonstrating that the large-scale topographic organization identified in neonates is robust to the inclusion or exclusion of the global signal. Warm colors indicate positive coupling, and cool colors indicate negative coupling between functional network topographies.

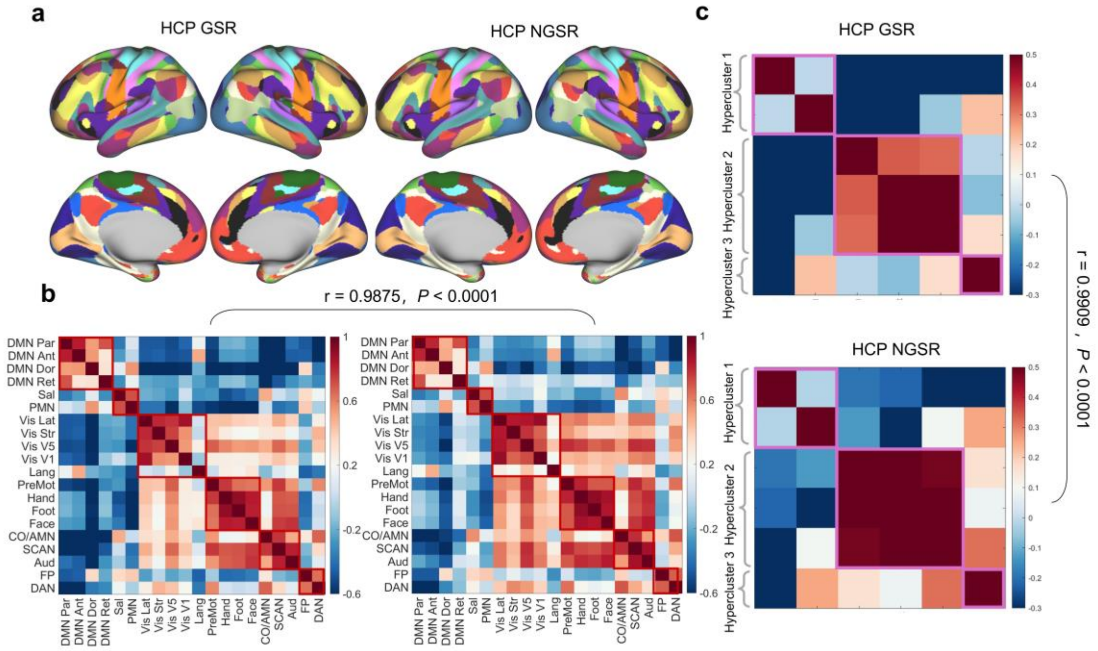


**Fig. S14 Robustness of adult functional network topography to global signal regression (GSR).** **a,** Group-level probabilistic topographies derived from the adult HCP dataset were converted into discrete network assignments by labeling each cortical vertex according to its highest network loading. Shown are group-level functional network parcellations for the HCP dataset computed with GSR (left) and without GSR (NGSR, right). **b,** The FOCA matrices from the adult HCP dataset were highly similar between the GSR and non-GSR conditions. **c,** The intercluster correlation matrices also showed high similarity with and without GSR, demonstrating that the large-scale topographic organization identified in the adult brain is robust to the inclusion or exclusion of the global signal. Warm colors indicate positive coupling, and cool colors indicate negative coupling between functional network topographies.

**Table S1 Similarity ($R^2$) of negative connectivity profiles between FOCA and FC across brain networks.**

| Network | DMN Par | DMN Ant | DMN Dor | DMN Ret | Vis Lat | Vis Str | Vis V5 | Vis V1 | FP | DAN |
|---|---|---|---|---|---|---|---|---|---|---|
| Similarity （$R^2$） | 0.92 | 0.73 | 0.49 | 0.83 | 0.86 | 0.86 | 0.94 | 0.73 | 0.87 | 0.95 |
| Network | PreMot | Lang | Sal | CO/AMN | PMN | Hand | Face | Foot | Aud | SCAN |
| Similarity （$R^2$） | 0.86 | 0.65 | 0.72 | 0.93 | 0.70 | 0.87 | 0.80 | 0.86 | 0.45 | 0.89 |

Note: DMN Par, DMN parietal; DMN Ant, DMN anterolateral; DMN Dor, DMN dorsolateral; DMN Ret, DMN retrosplenial; Vis Lat, visual lateral; Vis Str, visual streams; Vis V5, visual V5; Vis V1, visual V1; FP, frontoparietal; DAN, dorsal attention; PreMot, Premotor; Lan, language; Sal, salience; CO/AMN, cingulo-opercular/action-mode; PMN, parietal memory network; Aud, auditory; Hand, somatomotor-hand; Face, somatomotor-face; Foot, somatomotor-foot; SCAN, somato-cognitive-action network.
